# High-throughput DNA engineering by mating bacteria

**DOI:** 10.1101/2024.09.03.611066

**Authors:** Takeshi Matsui, Po-Hsiang Hung, Han Mei, Xianan Liu, Fangfei Li, John Collins, Weiyi Li, Darach Miller, Neil Wilson, Esteban Toro, Geoffrey J. Taghon, Gavin Sherlock, Sasha Levy

**Author notes:** Equal Contribution (Takeshi Matsui, Po-Hsiang Hung).

## Abstract

To reduce the operational friction and scale DNA engineering, we report here an *in vivo* DNA assembly technology platform called SCRIVENER (**S**equential **C**onjugation and **R**ecombination for **I**n **V**ivo **E**longation of **N**ucleotides with low **ER**rors). SCRIVENER combines bacterial conjugation, *in vivo* DNA cutting, and *in vivo* homologous recombination to seamlessly stitch blocks of DNA together by mating *E. coli* in large arrays or pools. This workflow is simpler, cheaper, and higher throughput than current DNA assembly approaches that require DNA to be moved in and out of cells at different procedural steps. We perform over 5,000 assemblies with two to 13 DNA blocks that range from 240 bp to 8 kb and show that SCRIVENER is capable of assembling constructs as long as 23 kb at relatively high throughput and fidelity. Most SCRIVENER errors are deletions between long interspersed repeats. However, SCRIVENER can overcome these errors by enabling assembly and sequence verification at high replication at a nominal additional cost per replicate. We show that SCRIVENER can be used to build combinatorial libraries in arrays or pools, and that DNA blocks onboarded into the platform can be repurposed and reused with any other DNA block in high throughput without a PCR step. Because of these features, DNA engineering with SCRIVENER has the potential to accelerate design-build-test-learn cycles of DNA products.

## Introduction

Scalable information processing platforms, such as those that handle written language or computer code, have sparked technology innovation cycles that develop applications to sit on top of these base layers. DNA holds promise to become the next major information medium, with emerging applications that use long DNA constructs to produce small molecules, multispecific antibodies, gene therapies, cellular therapies, and organisms designed to purpose.

However, DNA engineering is currently cumbersome, creating a bottleneck to the development of some DNA-based applications. For example, workflows for DNA assembly, such as restriction enzyme cloning, Gibson assembly^1^, and Golden Gate assembly^2,3^, can be idiosyncratic, expensive, and difficult to automate and scale. Clonal DNA must be purified from cells, manipulated using biochemistry and/or molecular biology approaches, and placed back into cells for cloning, amplification, and sequence verification. Scaling these *in vivo* → *in vitro* → *in vivo* DNA assembly workflows often requires construction of large centralized biofoundries with specialized equipment and experienced production teams.^4,5^

Thus, improving accessibility and reducing the operational friction of large scale DNA engineering has the potential to decentralize, reduce cost, and increase the pace of development of some DNA-based applications. To address this challenge, we introduce SCRIVENER (**S**equential **C**onjugation and **R**ecombination for **I**n **V**ivo **E**longation of **N**ucleotides with low **ER**rors), which combines elements of the MAGIC subcloning system^6^ with elements of the GENESIS and BASIS *E. coli*-based genome-scale assembly systems^7,8^ to enable assembly and sequence verification of long and difficult DNA constructs on plasmids at high throughput. SCRIVENER offers distinct advantages over existing DNA assembly and verification systems. Once DNA fragments are cloned into bacteria, the entire assembly process is as simple as mating and growing bacteria in 96- or 384-position formats. Arrays of uniquely barcoded recipient cell plasmids allow colonies to be pooled before DNA isolation and sequencing, significantly reducing the cost and time required for whole-plasmid sequence verification. Additionally, SCRIVENER streamlines the reuse of DNA fragments and assembled products, facilitating modular DNA assembly without PCR or other idiosyncratic *in vitro* manipulations.

## Results

### Overview of SCRIVENER

SCRIVENER sequentially and seamlessly assembles DNA blocks on plasmids simply by mating bacteria (**Fig. 1a**). DNA blocks are inserted into a fixed location on a ‘swapping cassette’ in donor plasmids and transformed into donor cells using standardized methods. Donor plasmids contain an origin of transfer (OriT) and are transferred via F-plasmid mediated conjugation from donor cells to recipient cells, which contain a recipient plasmid with a growing assembly next to another swapping cassette. A genetic program in the recipient cell cuts the swapping cassettes from both the donor and recipient plasmids using CRISPR/Cas9^9,10^ and “stitches” the incoming cassette into the recipient plasmid using the lambda Red homologous recombinase to extend an assembly^11^. Homology regions designed at the end of the incoming DNA blocks ensure that assemblies are seamless. Swapping cassettes feature alternating selectable and counter-selectable markers, facilitating iterative stitching. As in the MAGIC system and in contrast to BASIS and GENESIS, donor plasmids have a restricted origin of replication that is non-functional in recipient cells, ensuring they are lost during cell outgrowth and minimizing the possibility of unwanted recombination events. Recipient plasmid backbones contain a selectable marker (not shown in **Fig. 1a**), allowing for the selection of recipient cells and the elimination of donor cells. These design features ensure that most cells passing through the selection and counter-selection steps contain the desired product. DNA constructs are assembled on ColE1 plasmids, making it easy to extract sufficient DNA yield for downstream applications.

**Figure 1.**
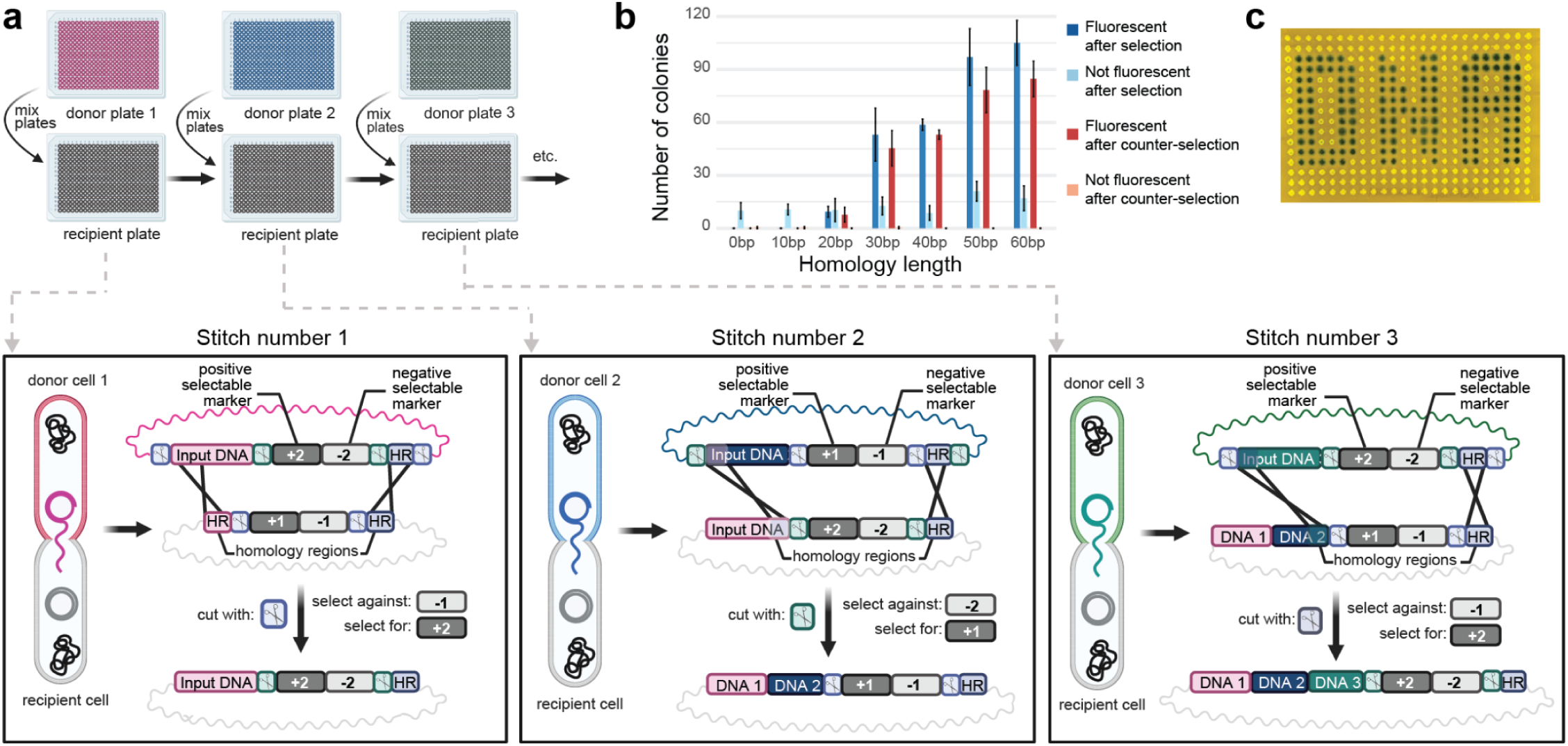
SCRIVENER schematic and fidelity. **a.** Arrays of input DNA blocks are introduced into donor plasmids and donor cells using standardized methods to create donor plates. In each assembly round, a DNA cassette is transferred from a donor plasmid to a recipient plasmid to extend a growing assembly. Two types of donor plasmids, used on alternating rounds, are transferred from donor cells to recipient cells via F-plasmid (not shown) mediated conjugation. Donor and recipient plasmids are cut in the recipient cell using CRISPR/Cas9 (scissors), guided by alternating gRNAs on the donor plasmids (not shown). Homology regions (HRs) promote recombination and seamlessly stitch input DNA together. Alternating selectable (+1, +2) and counter-selectable (−1, −2) markers on donor plasmids allow selection for sequential DNA transfers. **b.** Barplot showing the number of fluorescent (dark colors) and non-fluorescent (light colors) colonies following selection (blue) and counter-selection (red) for the last step in a liquid mPapaya assembly when the length of homology between the blocks is varied. Error bars show the standard deviation. **c.** Results from replicate four-part assemblies of mPapaya (fluorescent yellow colonies) and lacZ (blue colonies) grown on X-gal and visualized under blue light.

### DNA stitching fidelity, efficiency, and homology requirements

We first tested SCRIVENER by assembling the fluorophore mPapaya^12^ by sequentially stitching four DNA blocks (244 bases each) that contained 50-65 bp of overlapping homology. We successfully assembled the full *mPapaya* gene in liquid media and on agar, with all four independent replicates in both conditions being fluorescent and sequence perfect after the fourth stitch. To test how stitching accuracy and efficiency depends on the length of homology, we performed the final fluorescence-conferring stitch with DNA blocks that contained between 0 bp and 60 bp of homology (**Fig. 1b**). Fluorescent colonies were recoverable with as little as 20 bp of homology, but shorter homology lengths resulted in fewer colonies (reduced stitching efficiency) and a higher ratio of non-fluorescent colonies after selection (reduced stitching fidelity). After counter-selection, only fluorescent colonies remained in all conditions, indicating that counter-selection effectively removes incorrect recipient plasmids. Using this assay and cell counts over the course of the procedure, we estimated a stitching efficiency of ∼10^−5^ in liquid for stitching blocks with 50 bp of homology (**Supplementary Data 1**). For experiments that follow, we typically chose homology lengths of 45-50 bp, while avoiding sequence features that may lower homologous recombination efficiency and fidelity, such as repetitive sequences, secondary structure, homopolymers, and extreme GC content.

### DNA stitching and sequence verification in bacterial arrays

We next developed mid- and high-throughput protocols to multiplex SCRIVENER. For the mid-throughput protocol, we limited our toolset to those found in a standard molecular biology lab (96-well plates and multi-channel pipettes) with the aim of enabling most labs to perform hundreds of assemblies in parallel. For the high-throughput protocol, we assembled on 96- or 384-position arrays on agar with off-the-shelf pinning robots. We first tested both methods in a 96-position format with an expanded set of three fluorophores (mPapaya, mPlum, sfGFP, 4 stitches each) at 32-fold replication each. To test sequence fidelity in high throughput, we created arrays of recipient cells with unique DNA barcodes at each position. Assemblies on these pre-barcoded plasmids allowed us to pool arrays prior to DNA isolation and Oxford Nanopore Technologies (ONT) sequencing, significantly reducing the cost and time for whole-plasmid sequence verification.^13^ Using this sequencing method and a custom analysis pipeline, we found that 94 of 95 and 69 of 73 positions with sufficient sequencing coverage contained no errors using the liquid-based and agar-based protocols, respectively (**Methods**). We then tested a four-stitch assembly of the mPapaya and lacZ reporters in a 384-position format on agar. A functional reporter was observed at every position (**Fig. 1c**). These results demonstrate that many independent replicates of a construct can be assembled and sequence-verified at a nominal additional cost and time per replicate (**Supplementary Table 1**).

### SCRIVENER performance with long and challenging DNA sequences

Sequence features such as extreme GC content, homopolymers, tandem repeats, DNA secondary structure, long interspersed repeats, and long overall construct lengths^14^ are challenging to assemble by *in vitro* assembly methods that include PCR or DNA annealing steps. To test SCRIVENER performance across some of these features, we designed a test set of 21 constructs that includes natural Biosynthetic Gene Clusters (BGCs), regions of the yeast *Saccharomyces cerevisiae* genome (including a telomeric region and several regions that major DNA synthesis providers could not synthesize, **Supplementary Table 2**), and synthetic DNA constructs that contain different levels of overall GC content, DNA secondary structure, and interspersed repeats (**Fig. 2, Supplementary Data 2 and 3**). These assemblies used building blocks ranging from 1.8 kb to 5 kb to construct 8 kb to 23 kb constructs using 2 to 13 stitches at 12-fold replication each (**Fig. 2a**). To determine the assembly fidelity, we sequenced each assembly replicate either before (BGC and yeast genome assemblies) or after (synthetic DNA assemblies) isolation of a single cell at each position. Prior to single cell isolation, we expect each colony to be polyclonal, containing multiple competing cell lineages that stem from independent stitching events. A fraction of these lineages may contain detectable errors when sequenced at high coverage. After single cell isolation, we expect the colonies to be monoclonal. For each replicate, we calculated a purity score (**Extended Data Fig. 1**), representing the estimated fraction of plasmid molecules that are both full-length and have perfectly assembled sequences (**Methods**). We found that on average 66% of replicates per construct had a purity score >95% (**Fig. 2a**, **Supplementary Tables 3 and 4**).

**Figure 2.**
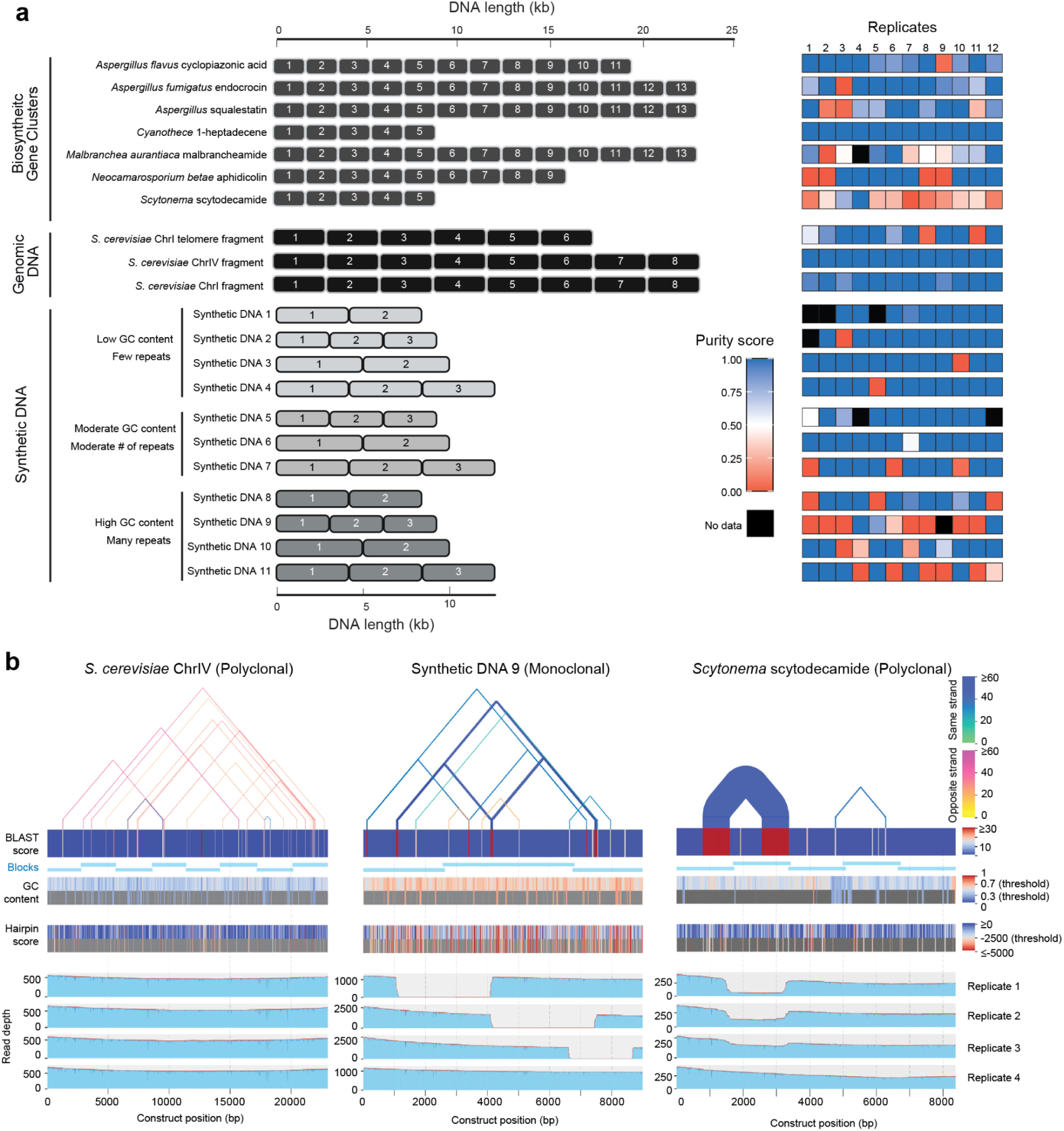
SCRIVENER performance with long and challenging DNA sequences. **a.** DNA blocks of different lengths (gray boxes) were assembled by sequential conjugation and selection on 96- or 384-position arrays at 12-fold replication. Plasmids in the resulting colonies were verified using high coverage long-read Oxford Nanopore sequencing. The purity score (heatmap) for each replicate indicates the fraction of sequence-perfect DNA molecules. **b.** Sequence feature (top) and read depth (bottom) plots of some replicates of long and/or complex assembled constructs. Regions that BLAST against each other (triangles) indicate long interspersed repeats, while those with high hairpin scores indicate DNA secondary structure. GC content is the GC% in a 51 bp rolling window. Regions with extreme GC% (<30% or >70%) and extreme secondary structure (hairpin score < −2,500) are highlighted in the lower half of each strip (gray box). Deletions between long interspersed repeats can be observed by a drop in read coverage in a replicate assembly.

### Characterization of SCRIVENER errors

For each construct, we examined sequence features (interspersed repeats, regions of extreme GC content, regions of secondary structure) that could contribute to errors along with the read depth coverage for each replicate (**Fig. 2b**). Inspection of these plots revealed that the most common error mode detected was a large deletion, typically between interspersed repeats on the same DNA strand (77 of 244 assemblies with sufficient sequencing coverage contained at least one deletion, ranging from 10 to 20,616 bp, **Supplementary Table 5**). The deleted regions typically encompassed a stitching junction and involved DNA from different stitching steps, suggesting that these deletions occurred after a successful stitch. Constructs with the lowest fidelity generally contained the longest imperfect direct repeats, and deletions frequently and reproducibly occurred between these repeats. For polyclonal colonies, we often found a mixture of full-length and deleted plasmid molecules (incomplete dips in the coverage at deletion loci in **Fig. 2b**), with the fraction of deletion-containing molecules being variable, even among replicates of the same construct. For putatively monoclonal colonies that went through a single cell isolation step, we generally found that all molecules either did or did not have a deletion. However, we detected some assembly replicates (13 of 125) that still contained mixtures of deleted and not-deleted molecules, indicating that multiple cell lineages were isolated, a single lineage contained multiple plasmids, or mutations occurred during cell outgrowth.

Examination of 50 bp windows flanking the predicted junction sites of the 117 detected deletions revealed a significant enrichment of long repeats in these regions (one-sided Wilcoxon rank-sum test p-value: 1.193×10^−38^, **Extended Data Fig. 2, Supplementary Table 5**), with 68 (59%) containing an imperfect repeat ≥20 bp (20-101 bp, average of 56.7 bp and 84% sequence identity). Other deletion junctions contained only shorter regions of more perfect homology (average of 12.2 bp and 90% sequence identity), suggesting Microhomology-Mediated End Joining (MMEJ) may be a mechanism in some cases.^15^ In addition, we found 4 cases (1.6% of all assemblies) where the deletions coincided exactly at the DNA block boundaries. This suggests that these are not deletions but partially assembled products from previous steps that counter-selection failed to eliminate during the stitching process. Another less common error mode detected was insertions (37 insertion events in 21 assemblies ranging from 11 to 8,342 bp, **Supplementary Table 6**), 43% of which were partial duplications of either the assembled product or the plasmid backbone, and 51% of which mapped to the *E. coli* genome or the F-plasmid. Notably, we did not observe any point mutations caused by SCRIVENER in the assembled product of full-length plasmid molecules, although some point mutations were found in the plasmid backbones. Point mutations were commonly detected in plasmid molecules with deletions or insertions, generally near the indel junctions (**Supplementary Table 5**).

Visual inspection of colonies during the assembly revealed that growth slowed for most assemblies when assembly lengths exceeded 15 kb and overall plasmid lengths exceeded 20 kb. This is presumably because these large relatively high-copy ColE1 plasmids pose a fitness cost on the cells that harbor them. We hypothesized that this fitness cost created a strong selective pressure to reduce the plasmid size. To test this hypothesis, we sequenced intermediates of the BGC assemblies at every odd stitch (stitch 1, 3, 5, etc.) and examined the assembly purity over time (**Fig. 3**). Read coverage plots at these intermediate stages revealed that deletions do not necessarily involve DNA actively being stitched. Instead, deletions can occur in DNA regions present early in the assembly, becoming apparent only when the overall plasmid size increases. Deletions typically expand in frequency over time suggesting that cells harboring these mutations are outcompeting cells with full-length products and eventually sweeping to dominate the cell population. The timing of the putative selective sweeps varied between replicates of the same assembly, suggesting that the rate at which the deletion mutation occurs and becomes established within the population is low, leading to stochastic, rather than deterministic, evolutionary dynamics^16^. The *Scytonema* scytodecamide BGC, which harbors an imperfect 823 bp interspersed repeat was an exception: most replicates harbored at least one lineage with a deletion at the repeat by the third stitch when the expected plasmid length is only ∼10.8 kb, suggesting that either the mutation rate to this deletion or the fitness advantage conferred is higher than other deletions in our BGC test set.

**Figure 3.**
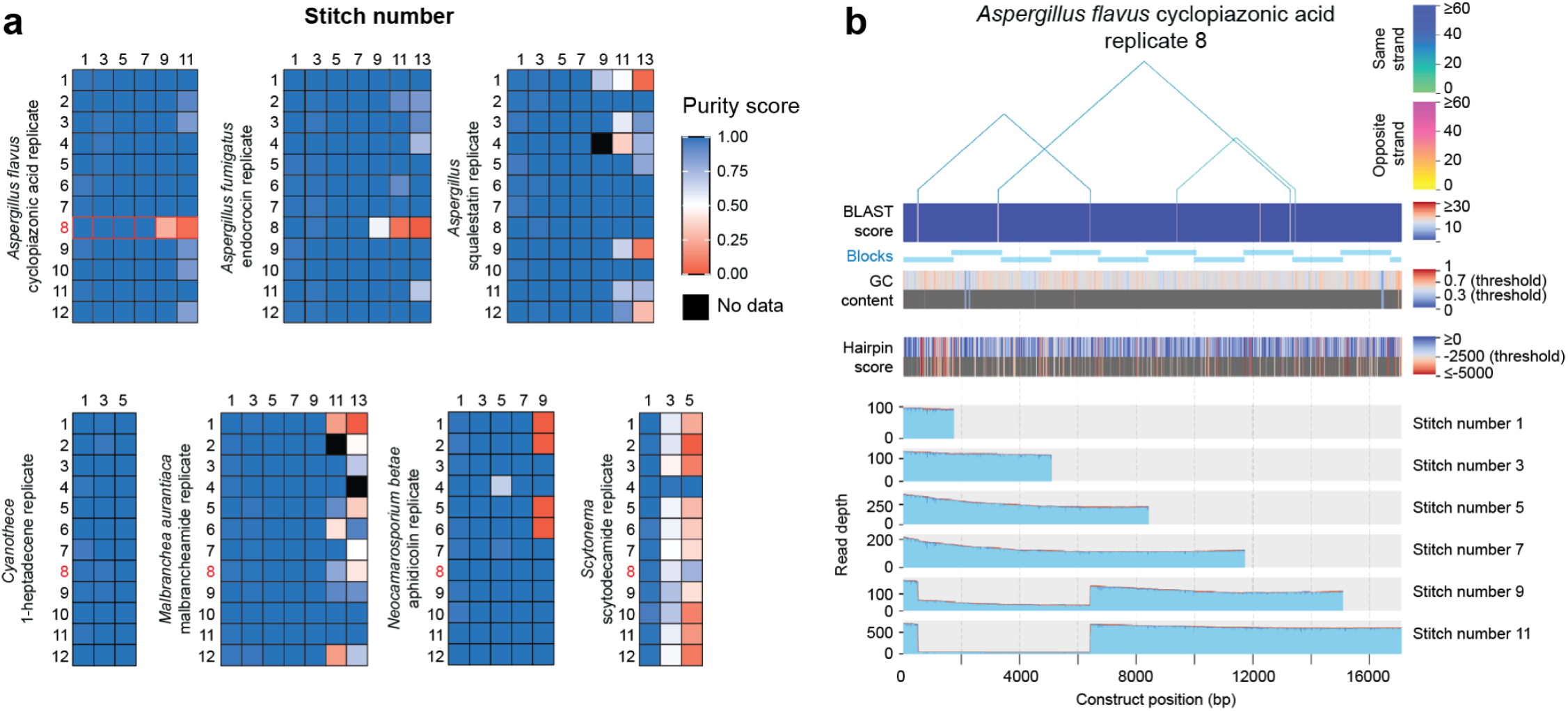
SCRIVENER deletion dynamics. **a.** Heatmaps of replicate assembly purity scores at different stages of assembly for biosynthetic gene clusters. Each row is a different assembly replicate, and each column is a different stage of assembly (stitch number). **b.** Sequence feature (top) and read depth (bottom) plots of a replicate of the *Aspergillus flavus* cyclopiazonic acid biosynthetic gene cluster (highlighted in red in panel **a**) at different stages of assembly.

### Construction of arrayed and pooled combinatorial libraries

To further demonstrate the ability for SCRIVENER to scale, we next attempted to construct an arrayed combinatorial library of 1,296 G-protein coupled receptor (GPCR) chimeras. We selected six human GPCRs (ADRB2, 5HT1A, 5HT1D, 5HT4R, AA2AR, AA2BR) and broke each into four blocks (181-618 bp), with fixed homology junctions between adjacent blocks positioned in transmembrane domains (**Fig. 4a, 4b**, **Supplementary Table 7**). We then assembled all 1,296 (6^4^) combinations at 4-fold replication on 384-position agar arrays. From these 5,184 assemblies, we observed growth of all colonies on the final stitch. Following sequencing, we detected sufficient reads from 4,777 (92%) of the positions representing 1,275 (98%) of the designs, at an average replication of 3.53 (**Supplementary Table 4**). Among these, 1,222 designs (94% of total) contained at least one replicate with a purity score >0.95, while 43 (3.6% of total) contained at least one replicate with sequence-perfect plasmid molecules but a lower purity score (2%-94.5%, 57.9% on average). Ten designs with sufficient sequencing coverage (0.77% of total) contained no sequence-perfect plasmid molecules in all replicates (purity score = 0), possibly because they contained cytotoxic DNA sequences.

**Figure 4.**
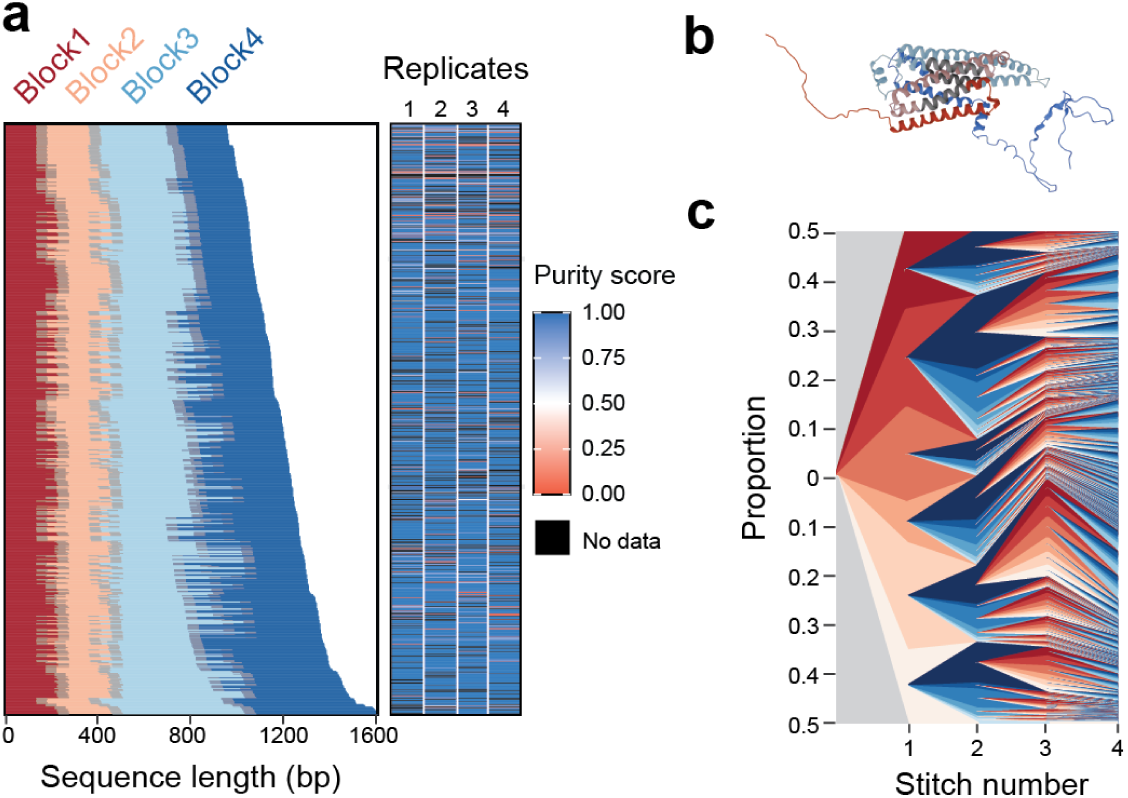
Assembly of combinatorial libraries in arrays and pools. **a.** Arrayed assembly of 1,296 combinatorial GPCR designs. Designs are ranked by overall length. Colors indicate the length of each block, with homologous overlaps shown in gray. Heatmap is the purity score for the four replicate assemblies of each design. **b.** Ribbon diagram of an assembled GPCR variant. Colors correspond to the blocks shown in panel **a**, with homologous overlaps shown in gray. **c**. Muller plot showing the relative frequency of each design over the course of a pooled assembly.

We next constructed the same combinatorial library using a simpler protocol whereby pools of donor and recipient bacteria are mated and undergo selection in 50 mL conical tubes (∼25,000 stitching events expected per cycle). We performed 4 independent pooled assemblies (2 sets of donor clones, 2 technical replicates each) and sequenced each pool at average coverage of 56, 74, 221 and 34 reads per design. We recovered 88%, 89%, 91%, and 70% of all 1,296 combinations, respectively, and detected 1,273 (98.2%) designs across the four pooled assembly replicates (**Fig. 4c**, **Extended Data Fig. 3, Supplementary Tables 8-10**). Examining the relative frequencies of each block in the pool revealed that some blocks were underrepresented and that frequency dispersion was likely to have a biological basis. Donor cells containing the same DNA block sometimes exhibited significantly and reproducibly different representation in the pool (**Extended Data Fig. 4** and 5). These results suggest that the stitching rate can vary among biological replicates containing the identical donor plasmid sequence, a finding that was not observable during assembly in arrays with a single donor clone at each position.

To estimate the stitching accuracy of pooled assembly, we isolated and sequenced 24 colonies from each pooled assembly replicate and found that on average 75% of the colonies from each pooled assembly replicate contained a sequence-perfect design. Most errors (18 out of 24 colonies, **Supplementary Table 11, Supplementary Data 4**) were partially assembled products from previous assembly steps, suggesting that the counter-selection condition we chose in these pooled experiments was insufficient to remove assembly midproducts.

### Reusing DNA blocks and assemblies without PCR

Composability, the ability to reuse and repurpose code snippets with low operational friction, has been a critical feature of information programming platforms because it accelerates the ability of engineers to build on top of each other’s work^17,18^. Repurposing of DNA blocks using existing *in vitro* methods generally requires that they are amplified with long primers by PCR to produce compatible ends, purified, validated, re-assembled, and transformed back into cells, a process that is difficult to automate and scale. By contrast, we reasoned that any two blocks already placed in SCRIVENER-compatible donor plasmids/cells could be seamlessly joined by utilizing small 100-140 bp “bridging” blocks that provide homology to both existing blocks. This workflow uses the same set of standardized operations as *de novo* DNA assembly, only some steps in the assembly process use bridging blocks. To test this idea, we attempted to join several of our previously incompatible blocks (no homology to each other) with bridges to their ends (**Fig. 5a, Supplementary Data 2 and 3**). As an added challenge, one of these assemblies contained an extremely long 2,519 bp imperfect repeat (88% sequence identity). Joining blocks with the same marker set required a single bridging block (with a different marker set), while joining blocks with different marker sets required two bridging steps. We found that all attempted bridges performed as designed, resulting in the expected chimeric assembly with high purity scores for most assembly replicates (**Supplementary Table 4**). However, the assemblies containing a 2,519 bp repeat had greatly reduced purity scores (0-28%), as expected from previous observations that deletions frequently occur between long repeats.

**Figure 5.**
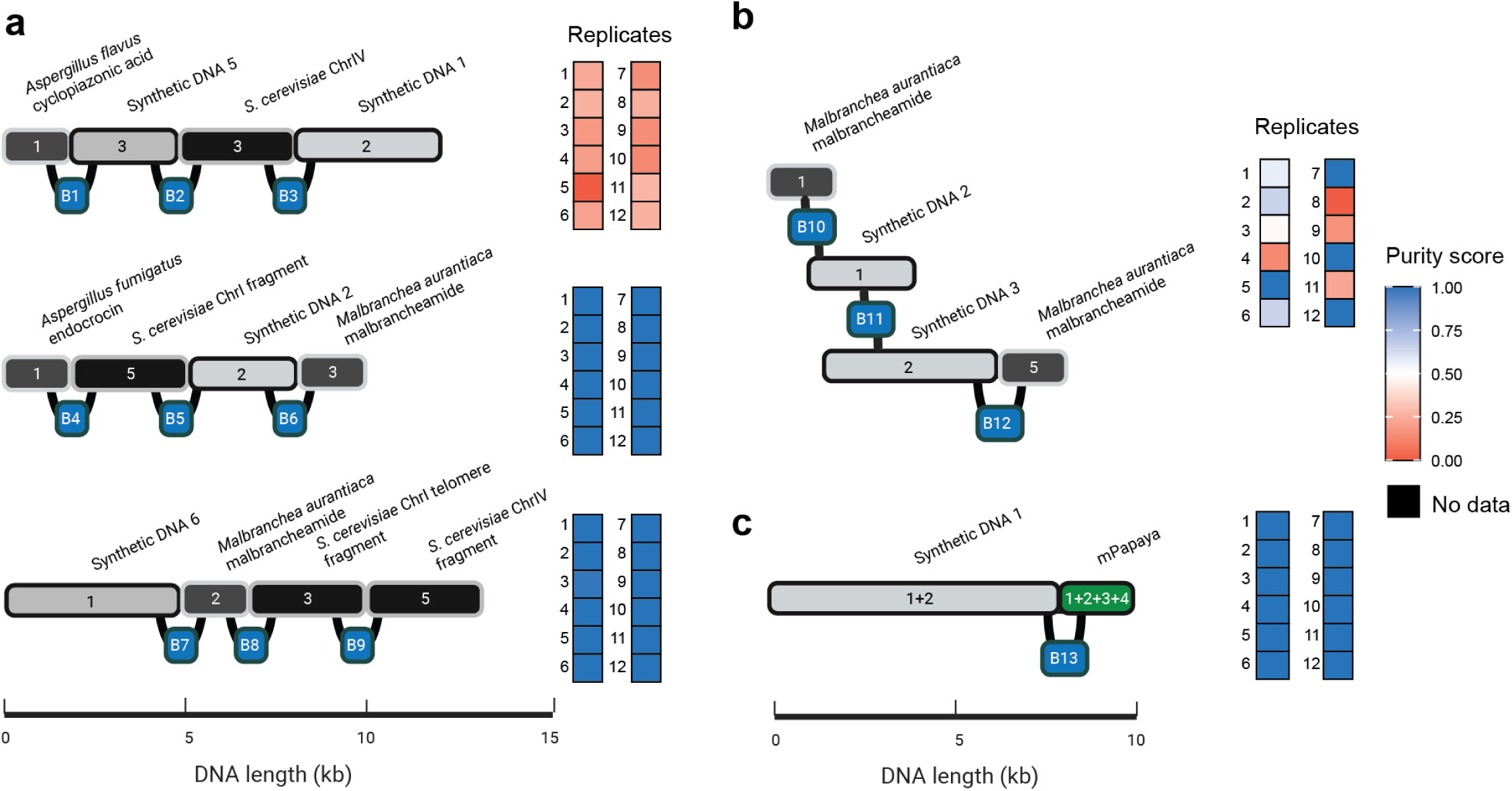
Composable DNA with bridging blocks. **a.** DNA blocks from different assemblies (numbered gray boxes) that lack homology to each other are seamlessly joined with 80-100 bp bridging blocks (B1-B9, blue boxes). The first assembly with low purity scores contains a 2,519 bp imperfect repeat between the synthetic DNA blocks, which is associated with frequent deletions. **b.** Bridging blocks (B10, B11) are designed with homology to the middle of an assembly or incoming block to join selected parts of existing blocks. **c.** Two complete assemblies are joined with a bridging block (B13). Heatmaps show the purity score for each replicate assembly.

We next asked whether a bridging block could be used to truncate an existing assembly or incoming block by providing homology to sequences in the middle, rather than the ends of these sequences. This would enable assembly of a chosen *part* of any existing block without PCR, thereby increasing the utility of DNA blocks already onboarded into SCRIVENER donors. We found that bridges with internal homology regions function as designed and can truncate at least 740 bp and 500 bp off of the end of a growing assembly and incoming block, respectively (**Fig. 5b and Supplementary Table 4**).

To test whether bridging blocks could be used to join fully assembled products, we transferred two previously assembled products from recipient plasmids to donor plasmids using standard restriction enzyme cloning. Our results showed that these products could be seamlessly joined with a bridging block (**Fig. 5c and Supplementary Table 4**), demonstrating the potential for hierarchical modular assembly of DNA constructs without PCR or other idiosyncratic *in vitro* manipulations.

## Discussion

We developed a high-throughput *in vivo* DNA assembly platform, SCRIVENER, that sequentially stitches DNA blocks together by mating bacteria in arrays or pools. SCRIVENER uses standardized protocols for onboarding of DNA blocks, mating, selection, and sequence verification, eliminating the need for idiosyncratic methods, expensive reagents (e.g., enzymes, kits), and procedures that require extended hands-on times (mixing reagents, running gels, quantitating DNA).

We show that SCRIVENER can assemble complex constructs up to 23 kb on commonly-used ColE1 vectors, but that long repeats and assembly lengths result in more frequent deletions, presumably because of selective pressure to relieve the fitness cost imposed by a large high-copy plasmid. Low-copy recipient vectors, such as the Bacterial Artificial Chromosomes (BACs) provide a promising avenue by which to extend lengths further^8^.

One limitation of SCRIVENER relative to Gibson and Golden Gate assembly is that SCRIVENER blocks must be assembled sequentially while *in vitro* methods can assemble several blocks in one step. However, the relative simplicity and low hands-on time of SCRIVENER enables more assemblies to be processed and sequence verified in parallel. Replicates add nominal additional cost and time to process, increasing the chances of recovering a sequence perfect clone on the first try for complex assemblies. Nevertheless, *in vitro* assembly strategies and SCRIVENER are likely to be highly complementary. For example, Golden Gate assembly could be used to insert multiple blocks into SCRIVENER donor plasmids, which are subsequently assembled *in vivo* to make larger constructs. We showed that complete assemblies can be ported back into donor plasmids and used to assemble longer constructs in a hierarchical workflow (**Fig. 5c**). While we used traditional *in vitro* cloning methods for this example, we envision that genetic programs similar to SCRIVENER stitching could be generated to perform this *DNA move* operation *in vivo*, potentially simplifying and increasing the speed of many-part assemblies.

In our hands, the greatest bottleneck to SCRIVENER throughput is onboarding DNA blocks into the donor plasmids/cells, even when using standardized methods. However, multiple DNA synthesis providers have scaled this process and offer DNA blocks already cloned into a vector of choice. Assuming these existing processes can be modified to function with SCRIVENER donor plasmids and cells, we envision that practitioners could purchase “stitch-ready” DNA blocks in donor plasmids/cells, enabling mid- to high-throughput DNA assembly simply by mating cells.

We demonstrated that SCRIVENER blocks are composable by using short bridging blocks. This feature could be useful for quickly generating new variants as part of design-build-test cycles, for constructing higher order assemblies that concatenate two or more small assemblies, and for shuffling the order of an assembly to optimize transcription or other properties. Bridging blocks can be designed to provide homology to the middle of a growing assembly or incoming block, thereby enabling selected parts of existing blocks to be reused in high throughput without a PCR step. While our initial tests indicated that bridges can truncate up to 740 bp off of a growing assembly and 500 bp off of an incoming block, further work is needed to assess practical truncation length limits. Depending on these results, it may be valuable to construct arrayed SCRIVENER donor cell libraries that create a reservoir of useful DNA sequences that can be reused without PCR (e.g., a human genome BAC library). Composability of the SCRIVENER platform could incentivize DNA block storage, reuse, and sharing, facilitating the development of a robust disaggregated DNA engineering ecosystem where practitioners can build on top of each other’s designs^19^. Instantiating such a DNA engineering environment requires products that further simplify SCRIVENER engineering (e.g. assembly kits and stitch-ready DNA blocks) and ecosystem infrastructure investments, such as those that increase the capacity of plasmid repositories or develop a SCRIVENER-compatible DNA block marketplace.

## Acknowledgements

We thank David Ross, Konrad Herbst, and David Craford for thoughtful comments on the manuscript. We thank Twist Biosciences, Inc. for providing some DNA blocks used in this study. We thank the NHGRI for their kind support of this work, including grants R01HG011676 to S.F.L and G.S. and R44HG012067 to S.F.L.

## Data availability

GenBank files of all plasmid backbones and table of DNA blocks used in this study are in the Supplementary Data. The raw FASTQ files from Oxford Nanopore sequencing conducted in this study are available in the Sequence Read Archive (SRA BioProject PRJNA1150152).

## Code availability

Code for sequence and data analysis is available in the GitHub repository https://github.com/tmatsui22222/SCRIVENER.

## Author contributions

X.L., T.M., W.L., D.M., and S.F.L conceived of the SCRIVENER system. T.M., P.H.H. and H.M. created the plasmids used in this study. H.M. engineered the *E. coli* strains. T.M., P.H.H., X.L., G.J.T., E.T., and S.F.L. designed experiments. T.M., P.H.H., H.M., and N.W. performed experiments. F.L and J.C. designed and wrote the sequence analysis pipeline. T.M., P.H.H., F.L., J.C., H.M., and D.M. analyzed data. T.M., P.H.H., and S.F.L. wrote the manuscript. All authors edited the manuscript. G.S. and S.F.L. provided suggestions for SCRIVENER design and experiments, and gave constructive feedback throughout the development of this work.

## Ethics declarations

Geoffrey J. Taghon was supported in part by an appointment to the NRC Research Associateship Program at the National Institute of Standards and Technology, administered by the Fellowships Office of the National Academies of Sciences, Engineering, and Medicine. Certain commercial equipment, instruments, or materials are identified to adequately specify the experimental procedure. Such identification implies neither recommendation or endorsement by the National Institute of Standards and Technology nor that the materials or equipment identified are necessarily the best available for the purpose. The views expressed in this publication are those of the authors and do not necessarily represent the views of the U.S. Department of Commerce or the National Institute of Standards and Technology.

## Extended Data Figures

**Extended Data Figure 1.**
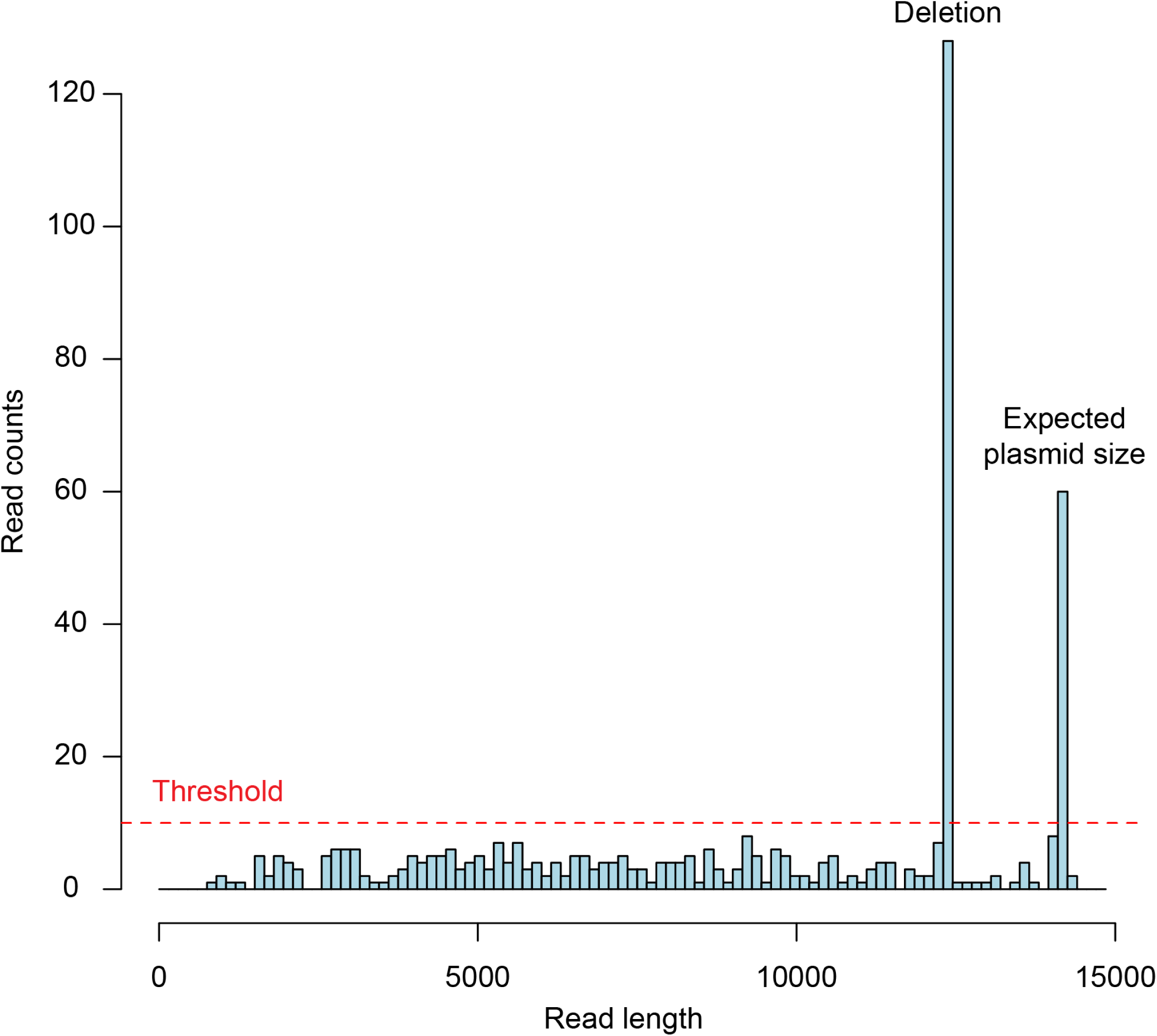
The histogram of read lengths for a replicate assembly of the *Scytonema* scytodecamide stitch number 5 shows two distinct “peaks”. These peaks presumably represent sequence data from fully intact plasmid DNA molecules that were not fragmented during plasmid extraction and library preparation, and suggests that there are at least two distinct plasmids within this replicate assembly. The red line is the read number threshold used to identify a peak. Any reads within 150 bp of these peaks were assigned to the closest peak to form “clusters”. In this example, two clusters with sizes of 12,373 bp (132 reads) and 14,173 bp (68 reads) were identified. The expected sequence length of this construct with the plasmid backbone is 14,174 bp, suggesting that one of the plasmids has a ∼1,800 bp deletion while the other plasmid is the expected size. To identify any sequence variants in these plasmids, reads in each cluster were analyzed individually using *bcftools* and *Sniffles2* (**Methods**). Using these data, purity score, defined as the fraction of plasmid molecules that are both full-length and have perfectly assembled sequences, was determined by calculating the fraction of reads assigned to the expected size cluster relative to the total number of reads assigned to all clusters. If any sequence variants were detected in the assembled product of the plasmid with the expected size, the purity score was set to 0. In this example, no sequence variants were detected in the plasmid with the expected size, and therefore, the purity score is 34% (68 / (132 + 68)).

**Extended Data Figure 2.**
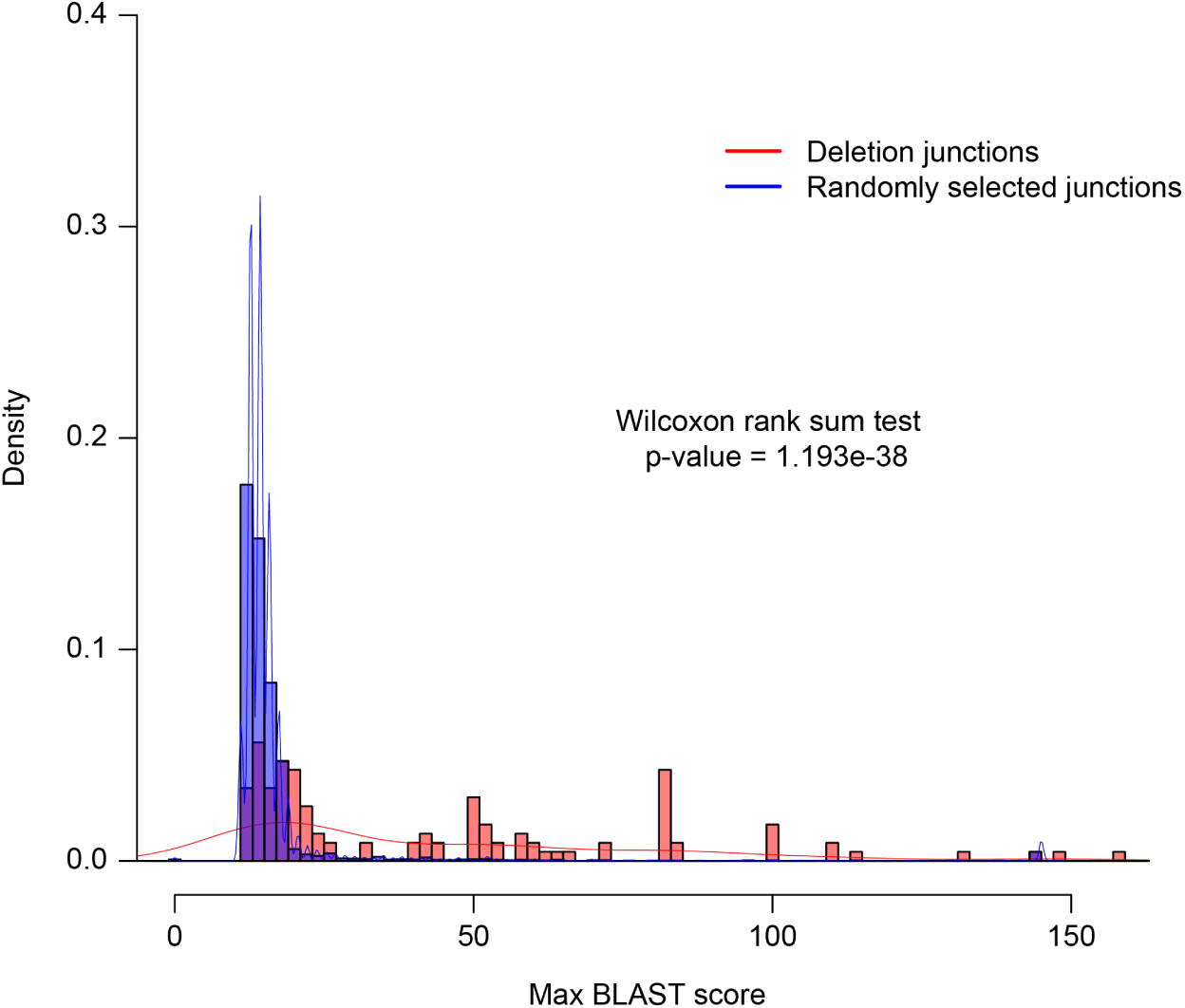
Density plots comparing the maximum blast score of 117 deletion junctions found in the long and difficult DNA test set (red) with random junctions (blue). The random junctions are separated by the same distance as the deletions and were chosen from the same construct where the deletion was detected.

**Extended Data Figure 3.**
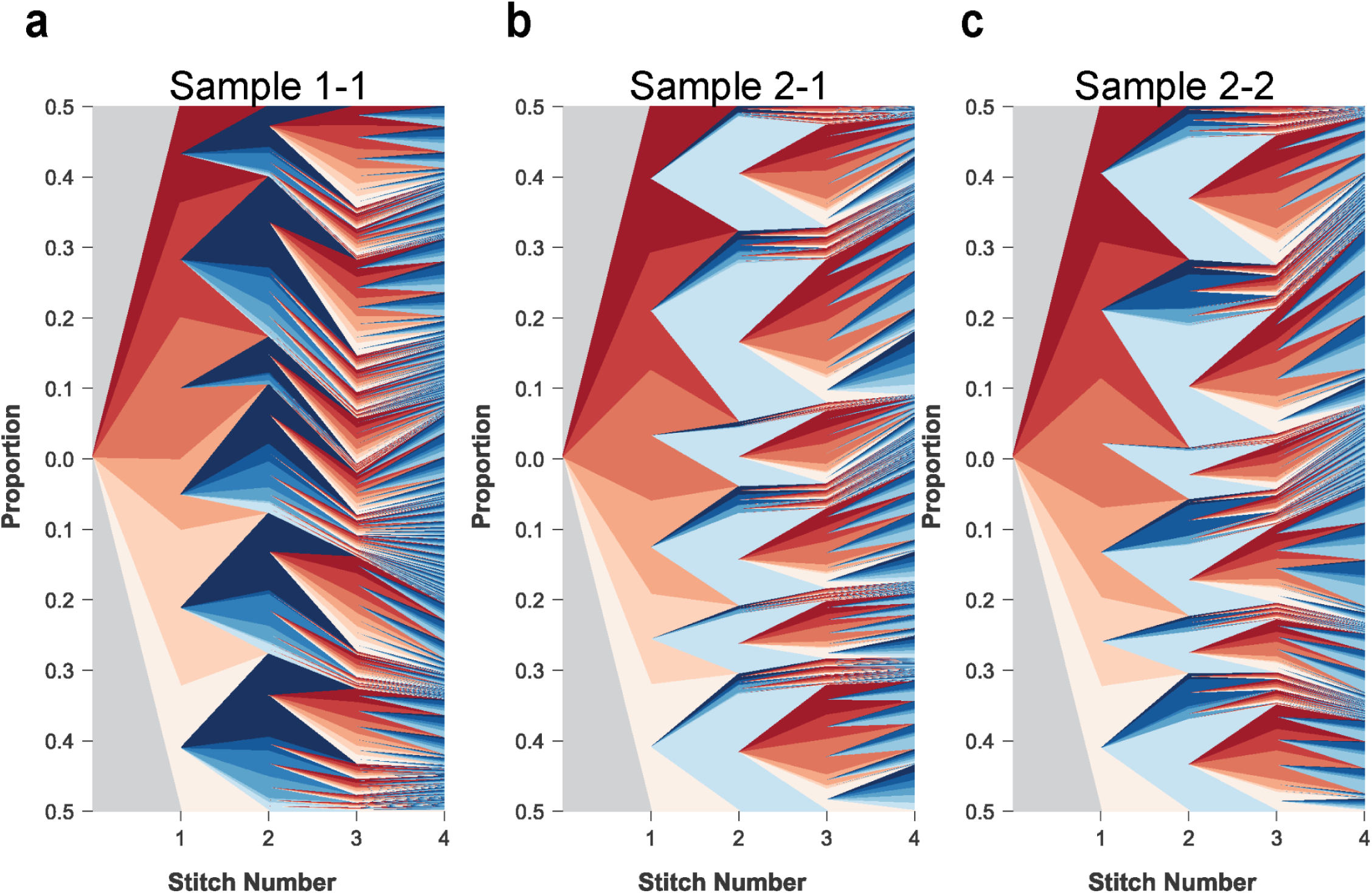
Muller plots showing the frequency distribution of each assembly product at every stitch for pool assembly replicates 1-1 (**a**), 2-1 (**b**), and 2-2 (**c**). Replicate 1-2 is shown in Fig. 4C.

**Extended Data Figure 4.**
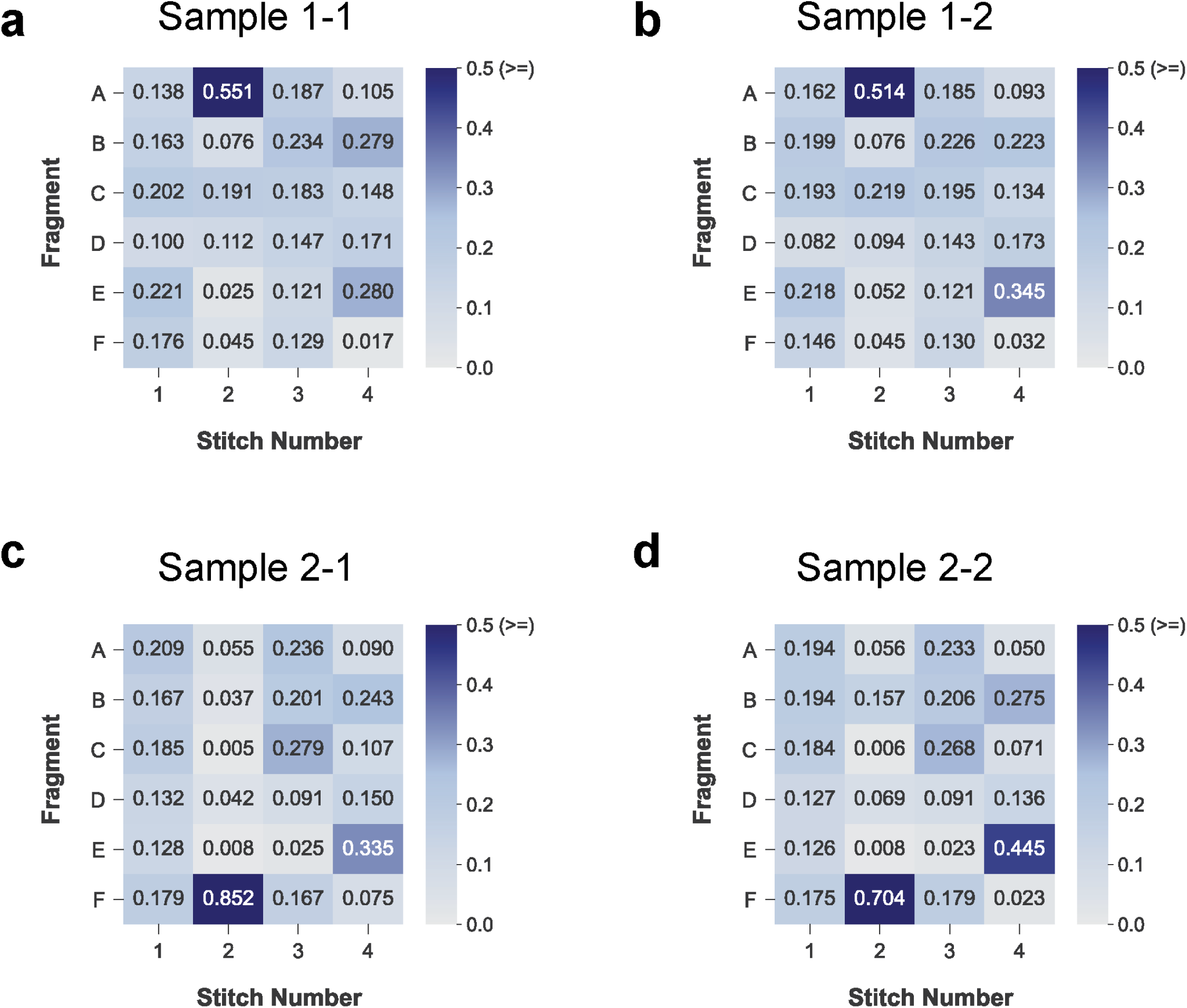
The frequency of each DNA block in the final pool for each pooled assembly replicate. Colors represent the relative frequency of each block. The expected value is 0.167 (1/6) for each block.

**Extended Data Figure 5.**
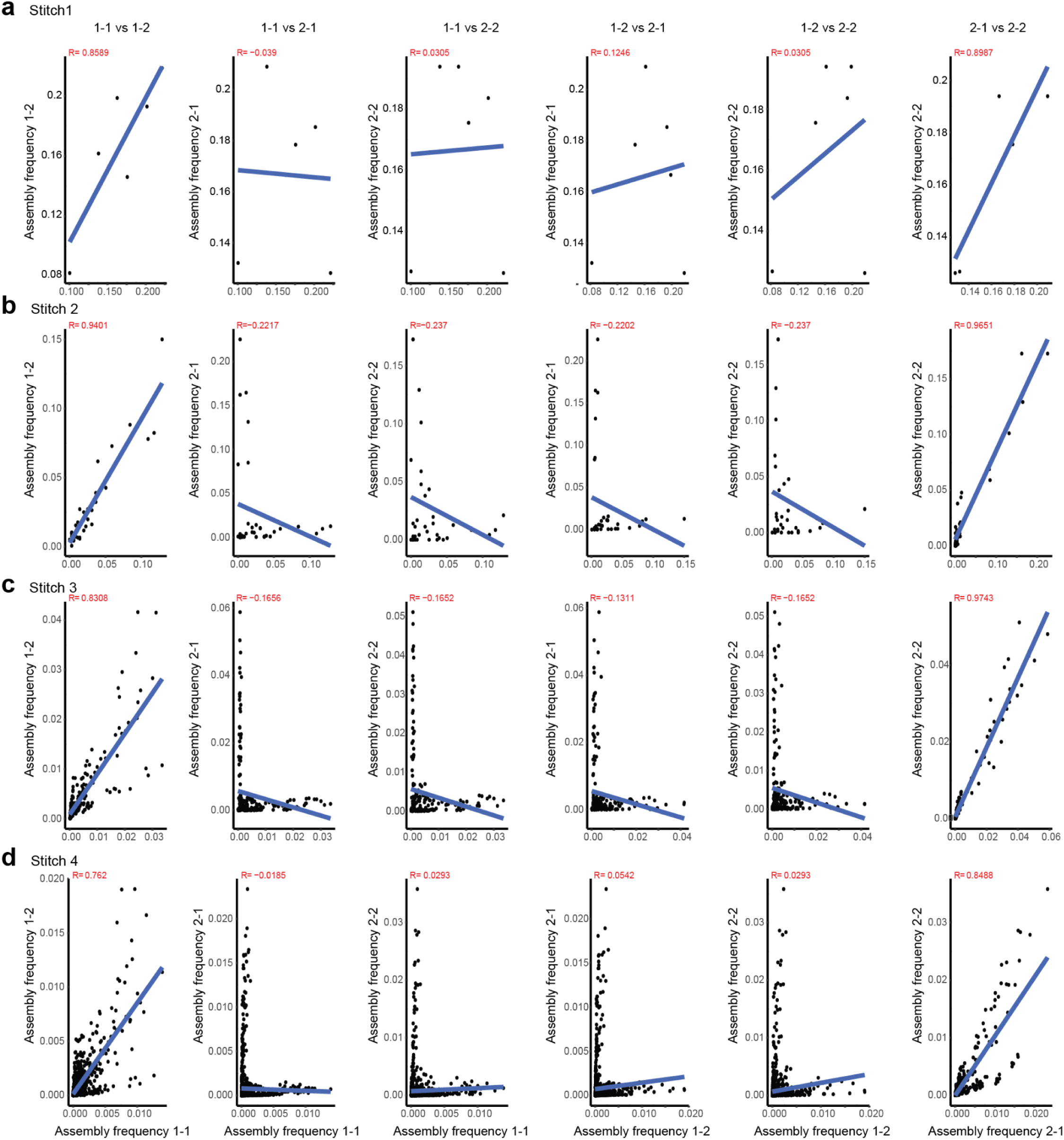
Scatter plot of the relative read frequency of each pooled assembly replicate across four stitches: (**a**) Stitch 1, (**b**), Stitch 2, (**c**) Stitch 3, and (**d**) Stitch 4. Technical replicates are 1-1 vs. 1-2 and 2-1 vs. 2-2, and biological replicates are 1-1 vs. 2-1, 1-1 vs. 2-2, 1-2 vs. 2-1, and 1-2 vs. 2-2. Blue line and red text indicate the Pearson correlation (R).

## Supplementary Data

**Supplementary Data 1.** Results from replicate experiments used to calculate the stitching efficiency. **Tab 1 (CFU_per_ml):** Results from replicate experiments used to calculate Colony Forming Units (CFUs) for donor cells carrying the fourth block of a mPapaya assembly and recipient cells carrying the first three blocks. These cells were used to assay the stitching efficiency with 50 bp of homology. **Tab 2 (stitching_efficiency)** Results from mating donor and recipient cells, used to calculate the stitching efficiency. **Tab3 (colonies_pass_selection)** Results from mating donors carrying the fourth block of mPapaya with different homology lengths (0-60 bp) with the third mPapaya assembly recipient cells after selection. **Tab4 (colonies_pass_counter-selection)** Results from mating donors carrying the fourth block of mPapaya with different homology lengths (0-60 bp) with the third mPapaya assembly recipient cells after selection and counter-selection.

**Supplementary Data 2.** The sequences of the DNA blocks (**Tab 1: DNA blocks**) and bridging blocks (**Tab 2: bridge**). GPCR blocks from *ADRB2*, *5HT1A*, *5HT1D*, *5HT4R*, *AA2AR*, and *AA2BR*, are denoted as A, B, C, D, E, and F, respectively. GPCR design names are named based on the specific GPCR DNA blocks used in each stitch. For instance, a GPCR design named ABCD indicates that the first DNA block is from *ADRB2*, the second is from *5HT1A*, the third is from *5HT1D*, and the fourth is from *5HT4R*.

**Supplementary Data 3.** The sequence of each construct assembled in this study is provided, along with the corresponding expected plasmid sequence including the plasmid backbone. The 20 bp stretches of Ns in the expected plasmid sequence represent the locations of the recipient plasmid barcodes. The expected plasmid sequences are organized with the construct sequence beginning at position 1, followed by the plasmid backbone sequence.

**Supplementary Data 4.** Fasta files containing the consensus plasmid sequences from single colony isolates of the pooled GPCR assemblies.

**Supplementary Data 5.** Genbank files of plasmids used in this study.

## Supplementary Tables

**Supplementary Table 1.** Estimated costs for different steps in an *in vivo* DNA assembly workflow. SCRIVENER HT is assembly on agar. SCRIVENER MT is assembly in liquid media in 96-well plates.

**Supplementary Table 2.** DNA blocks used to construct yeast genomic DNA assemblies were tested for synthesizability by Twist and IDT through an API. A summary of the results from both vendors is shown.

**Supplementary Table 3.** Summary table of purity score results for the 244 replicate assemblies from the long and difficult DNA test set. “Mean_purity” column is the average purity score across all replicates with sufficient sequence coverage for that construct. “Number_of_correct_replicates” indicates the number of replicates with sequence coverage of at least 20 and a purity score >0.95.

**Supplementary Table 4.** Assembly results for all replicate assemblies in this study are presented as follows: The “Well” column indicates the position on the 96- or 384-well plate where the construct was assembled, with each position representing an independent assembly replicate for a construct. The “Ref_size” column denotes the expected size of the assembled product in base pairs (bp). The “Peaks” column lists the read lengths of detected peaks in the read length distribution using our custom analysis pipeline (**Extended Data Fig. 1**). The “Clustered_coverage” column shows the total number of reads assigned to a cluster. The “Clonality_estimate” column provides an estimate of the number of different plasmid molecules present within an assembly replicate, based on the number of detected peaks. “Purity_score” is the estimated fraction of sequence perfect full length plasmid molecules within a replicate assembly. Any rows with a purity score of “NA” indicates that the sequence coverage was too low to confidently estimate the purity score. The “Assembled_product_mutation” and “Backbone_mutation” columns indicate whether any mutations were detected in the assembled product and plasmid backbone, respectively, for reads in the expected size cluster. The “Assembly_mutation” column specifies whether a mutation in the assembled product was introduced during assembly (“TRUE”) or was already present in the donor plasmid (“FALSE”). The “Native_barcode” column identifies the rapid barcode used for multiplexing the samples during library preparation, while the “BC_sequence” column provides the 20mer barcode present in the recipient plasmid. Lastly, the “Raw_data_file” column is the name of the file on SRA containing the raw fastq reads for each assembly replicate.

**Supplementary Table 5.** Sequence variants identified in this study are detailed as follows: The “Start” and “End” columns represent the start and end positions of the observed variants, respectively. The “Variant_type” column categorizes the variants into point mutation, deletion, insertion, or inversion. The “Variant_length” column specifies the length of the variants in base pairs (bp). The “Reference” column presents the sequences in the reference at the position where the variant was identified, while the “Alt” column shows the observed sequence(s). The “Quality_score” indicates the confidence level of the variant call, with higher values signifying greater confidence. Different quality score scales are used for *bcftools* and *Sniffles2.* For point mutations and small indel calls made with *bcftools,* only mutations with a quality score of above 100 are reported. For large insertions, deletions, and inversions called by *Sniffles2,* only variant calls with a quality score of above 20 are reported. The “Location” and “Cluster_type” columns specify the location of the variant (either in the assembled product or the plasmid backbone) and the read cluster in which the variant was identified (either in the expected size cluster or a cluster of the wrong size), respectively.

**Supplementary Table 6.** Alignment results for 37 distinct insertion events observed in the challenging DNA test set. “Insert_length” column is the length of the insertion in base pairs, while the “match_length” column is the number of base pairs within the insertion that matched to one of the reference plasmid sequences. The “Percent_Mapped” column shows the percentage of alignment between the sequence from “match_length” and the reference. The “reference_match” column identifies the reference to which the insertion sequence had the best match.

**Supplementary Table 7.** Alignment of amino acid sequences of the 6 human GPCRs assembled in this study. The 6 GPCRs are structure-based alignment by GPCRdb^20^. GPCR blocks from *ADRB2*, *5HT1A*, *5HT1D*, *5HT4R*, *AA2AR*, and *AA2BR*, are denoted as A, B, C, D, E, and F, respectively. The “Blocks” row shows the 4 blocks used for SCRIVENER assemblies: Block 1 (dark red), Block 2 (light red), Block 3 (light blue), and Block 4 (dark blue). Linker sequences (45 bp), shown in gray, and are contained within both adjacent blocks to facilitate homologous recombination.

**Supplementary Table 8.** Read coverage and number of GPCR designs detected at various steps of filtering for the pooled GPCR library assembly data. Column 2, “Stitch#”, indicates the number of stitching steps. Column 3, “Repeats,” specifies the sample source, where the first number represents the biological replicate and the second number represents the technical replicate. For example, 1-1 and 1-2 are technical replicates where the same donor clones were used for stitching. Column 4, “Total reads”, shows the number of raw reads obtained from nanopore sequencing. Column 5, “Total mapped reads”, represents the reads that align with the reference GPCR variants using minimap2. Column 6, “Total unique mapped reads”, is the count after filtering the minimap2 results, where each read is assigned to only one GPCR variant based on the matched alignment length. Reads matching multiple GPCR variants with the same alignment length are excluded from the analysis. Column 7, “# of variants expected”, indicates the expected number of GPCR variants at each step. Column 8, “Average coverage”, is calculated as the ratio of Column 6 to Column 7, representing the average coverage of a GPCR. Column 9, “Identified variants”, denotes the unique GPCR variants identified from each pool based on the total unique mapped reads. Column 10, “Successful rate”, is the ratio of identified unique GPCR variants (Column 9) to the expected number of GPCR variants (Column 7). Column 11, “Missed variants”, lists the variants not detected, with some entries including two numbers. The numbers in parentheses are corrected counts. Some variants may exist in the pool but were undetected initially; their presence is inferred if their descendant GPCR variants are detected in subsequent stitches, indicating the parental GPCRs were present but not detected. Also see Table S9 for details.

**Supplementary Table 9.** Table of missed GPCR variants. This table details all the missed GPCRs at each step and is used to correct the actual assembled GPCRs at each step in Table S8. The parental GPCR variant refers to the GPCR variant from the previous stitch (Column 3). Each parental GPCR variant can generate 6 expected child GPCR variants. If all 6 child GPCRs are missing, we assume the parental GPCR was not in the pool on the previous stitch. However, if any child GPCR is present, we assume the parental GPCR was also present in the previous stitch.

**Supplementary Table 10.** Data analysis summary for the pooled GPCR assembly. Column 1, “unique ID”, is a unique ID for each GPCR variant. Column 2, “GPCR”, denotes the GPCR variant assembled at each step. “R” represents the recipient strain, which is always the first parental strain used for stitching. Column 3, “Parent” is the unique id (Column 1) of the parent from the previous stitching step. Column 4, “#Stitch”, indicates the stitch number. Columns 5-8 are the added GPCR blocks in each stitching step. Columns 9-12, “R_X-X”, show the number of sequencing reads that matched to the indicated GPCRs in each replicate. The first number in X-X represents the biological replicates, and the second number represents the technical replicates. For example, 1-1 and 1-2 are technical replicates where the same donor clones were used for stitching.

**Supplementary Table 11.** Results from sequencing of single clones isolated from pooled GPCR assemblies. The “GPCR_pooled_replicate” column indicates the pooled GPCR assembly replicate from which the single colonies were isolated, while the “replicate” column specifies the individual clonal isolate number (24 for each pooled GPCR assembly replicate). The “Query_length” column denotes the length of the plasmid isolated from each single isolate in base pairs. “Query_start” and “query_end” represent the start and end positions on the plasmid where the sequence matches one of the GPCR designs. The “Target”, “target_length”, “target_start”, “target_end”, and “residue_match” columns display the GPCR designs to which the single isolate sequence best matched, along with the start and end positions of the match and the number of exact residue matches. The “Match_status” column indicates whether the match was a full-length exact match to one of the GPCR designs (“Match”), contained some mutations (“Mutation”), was an assembly product from previous assembly steps (“Mid-product”), or did not match any GPCR designs (“No-match”).

**Supplementary Table 12.** Plasmids used in this study

**Supplementary Table 13.** Primers used in this study.

## Materials and Methods

### Strains used in this study

The donor strain (dSL2: HB101 *ΔuidA::pir+*, *ΔendA::FRT, F128-xx (oriT::TcR)*) and recipient strain (rSL2: HB101 *ΔendA::FRT*) were constructed from the HB101 strain^21^ (*araC14, leuB6(Am), Δ(gpt-proA)62, lacY1, glnX44(AS), galK2(Oc), λ-, recA13, rpsL20(strR), xylA5, mtl-1, thiE1, [hsdS20]*) using a pop-in pop-out strategy, as described^22^. The *endA* gene was deleted in both donor and recipient strains to increase plasmid stability. Insertion of *pir+ gene* in the genome of the donor strain was necessary to support replication of the donor plasmids with an R6Kγ origin of replication^23^. The *oriT* in the F128-xx plasmid was replaced with a tetracycline resistance marker (TcR) to prevent conjugation of the F plasmid itself to the recipient strain.

### Media and chemicals

Luria-Bertani (LB) broth was used for cloning and for growth of donor and recipient plasmids. For selection and maintenance plasmids, antibiotics were added at the following concentrations: kanamycin (Kan) (25 *μ*g/mL), spectinomycin (Sp) (60 *μ*g/mL), hygromycin B (Hyg) (100 *μ*g/mL), gentamicin (Gm) (25 *μ*g/mL), tetracycline (Tc) (2 *μ*g/mL), carbenicillin (Carb) (100 *μ*g/mL), chloramphenicol (Cm) (25 *μ*g/mL), apramycin (Apm) (100 *μ*g/mL), streptomycin (Str) (25 *μ*g/mL). For counter-selection and removal of plasmids, 6% sucrose was added for *SacB*, and 200 *μ*g/mL 4-chloro-phenylalanine (4CP) was added for *PheS*. To improve the efficiency of *PheS* counterselection in LB, T251A/ A294G *ePheS* variant was used^24^. L-arabinose (0.2% w/v) and L-rhamnose (0.2% w/v) were used to induce the *ParaBAD* and *PrhaBAD* promoters, respectively.

### Plasmids used in this study

All plasmid backbones used in this study were constructed by restriction digestion and ligation or Gibson assembly of synthetic gene blocks sourced from Twist, IDT, or GenScript and are listed in **Supplementary Table 12** and **Supplementary Data 5**.

### Construction of barcoded recipient cell arrays

To generate arrays of recipient strains with uniquely barcoded recipient plasmid, the rSL2 recipient strain was transformed with helper plasmid eBSD28 and selected for integration using LB+100 *μ*g/mL Carb to create eBSD35. The helper plasmid eBSD28 contains an arabinose inducible *Cas9*, rhamnose inducible lambda red recombineering genes, and a temperature-sensitive *pSC101* origin. Barcoded recipient plasmids were constructed by inserting a random 20mer into the eBSD10 plasmid backbone region. To achieve this, eBSD10 was first PCR amplified using 2 pairs of primers: oSL1482 (which contains the 20mer barcode) and oSL1484, and oSL1426 and oSL1483 (**Supplementary Table 13**). The PCR amplified products were gel extracted and assembled via NEBuilder HiFi DNA assembly master mix (NEB). The assembled product was transformed into eBSD35, selected on LB+25 *μ*g/mL Gm, and 672 single colonies were picked. The barcode region from presumptive clones was amplified using primers oBSD20 and oBSD9 and Sanger sequenced. Unique barcodes with a Levenshtein distance greater than 6 to each other were selected and rearrayed to generate three separate arrays of barcoded recipient clones (paSL5 - 96 barcodes, paSL6 - 384 barcodes, and eaBSD3 - 96 barcodes, 576 barcodes total).

### Insertion of DNA blocks into donor plasmids

DNA blocks used for assemblies were synthesized by Twist Bioscience (**Supplementary Data 2**), IDT, or amplified from BY4716 yeast genomic DNA by PCR via PrimeSTAR or Platinum SuperFi II DNA Polymerase (Thermo Fisher Scientific) (**Supplementary Table 13**). The blocks were cloned into linearized donor vectors by standardized methods using either restriction enzyme cloning, Gibson cloning, or gap repair cloning. For restriction enzyme cloning, the DNA block and the appropriate donor plasmid were digested by the restriction enzymes AscI and NotI and purified by gel extraction. The purified plasmid backbone and insert was ligated by T4 ligase reaction following the standard protocol (NEB). The ligated products were transformed into the dSL2 donor cell and selected on the appropriate LB+antibiotic plates.

For Gibson cloning, the appropriate linearized plasmid and purified DNA block was mixed into the NEBuilder HiFi DNA assembly master mix following standard protocol (NEB) and then transformed into dSL2 donor cells. Overlaps of 20-25 bp were used.

For gap repair cloning, 200 ng of the DNA block and 200 ng of the appropriate linearized donor plasmid were transformed into dSL2 donor cells to assemble the two fragments via the endogenous E. coli machinery^25^. Overlaps of 20-25 bp were used.

All cloned plasmids were purified using a Qiagen miniprep. For small inserts, insertion regions were verified by Sanger sequencing (McLAB or Azenta/Genewiz). For larger inserts, whole plasmids were sequenced by Oxford Nanopore sequencing (Plasmidsaurus, Primordium, or in-house).

### Testing DNA stitching fidelity, efficiency, and homology requirements for SCRIVENER

The mPapaya fluorescence gene was split into four DNA blocks such that four stitches are necessary to complete the assembly (**Supplementary Data 2**). All blocks were inserted into the appropriate donor plasmid/strain (eBSD13, eBSD16, or eBSD19 in strain dSL2) The first three DNA blocks were stitched into recipient strain rSL2 and tests were performed by stitching the fourth DNA block. Several mPapaya fourth donors were designed such that the donor blocks have 0, 10, 20, 30, 40, 50, and 60 bp of homology with the third mPapaya stitch product. The recipient strains and the different donor strains were first grown overnight in LB+60 *μ*g/mL Sp+25 *μ*g/mL Gm+100 *μ*g/mL Carb and LB+100 *μ*g/mL Hyg+2 *μ*g/mL Tc, respectively. The cells were then diluted 100 fold with the same LB+antibiotic and grown at 30°C for four hours. After growth, OD600 was measured for each culture, and each culture was then normalized to OD600 of 0.5. Twenty-five *μ*L of the recipient strain was mixed with 75 *μ*L of one of the donor strains, with three replicates for each recipient-donor pair. The cell mixtures were centrifuged and re-suspended in 1 mL of SOB+0.2% arabinose+0.2% rhamnose and grown at 30°C for two hours. After growth, OD600 was measured again for each culture to make sure a similar number of cells were being plated after mating across all conditions. Fifty *μ*L of the culture was plated onto LB+50 *μ*g/mL Sp+25 *μ*g/mL Gm+100 *μ*g/mL Carb agar plates and grown for two days at 30°C. The cells were then replica-plated using a velvet onto LB+50 *μ*g/mL Sp+25 *μ*g/mL Gm+100 *μ*g/mL Carb+6% sucrose agar plates and grown at 30°C for one day. The total number of colonies and the number of fluorescent colonies was counted for each and used for calculation of relative stitching fidelity and efficiency (**Supplementary Data 1**).

To estimate the absolute stitching efficiency for the stitch with 50 bp of homology, we first determined the colony-forming units (CFU) in a 1 mL culture at an OD600 of 1 for both the recipient and donor strains. A recipient strain carrying the third mPapaya stitch product and a donor carrying the fourth mPapaya DNA block with 50 bp homology were grown overnight in LB+60 *μ*g/mL Sp+25 *μ*g/mL Gm+100 *μ*g/mL Carb and LB+100 *μ*g/mL Hyg, respectively. They were then diluted 100 fold and grown at 30°C for four hours. After growth, OD600 was measured for each replicate and each culture was normalized to either OD600 of 0.25, 0.5, and 0.75, with three replicates each. The normalized cultures were further diluted to 10^−5^ in 1 mL of SOB, and 100 *μ*L of this dilution (for a total of 10^−6^ dilution) was plated onto either LB+60 *μ*g/mL Sp+25 *μ*g/mL Gm+100 *μ*g/mL Carb agar plate for recipient strains and LB+100 *μ*g/mL Hyg agar plates for donor strains. The cells were grown for two days at 30°C and the total number of cells was counted (**Supplementary Data 1**). To get the CFUs per 1 mL when OD600 equals 1, the number of colonies was multiplied by 10^6^ *(1/initial OD600). On average, a CFU was ∼1.5 × 10^8^/mL for donor strains and ∼3 × 10^8^/mL for recipient strains. Using this CFU value for the recipients, stitching efficiency was determined according to the following equation: efficiency = ((number of colonies after counter-selection)) / ((OD600 before mating / OD600 after mating) x (recipient CFU per 1 mL when OD600 equals to 1) x (volume used for mating in *μ*L/1,000 *μ*L/mL)). The efficiency measured for each replicate was 9.05 × 10^−6^, 1.23 × 10^−5^, and 1.04 × 10^−5^, for an average efficiency of 1.06 × 10^−5^. These estimates are conservative because they assume that all stitching happens immediately upon mating and all outgrowth following mating is creating replicates of successful stitching events. In reality, some outgrowth is happening before stitching, which, if measurable and taken into consideration, would increase the efficiency we calculated.

### Arrayed assembly on agar plates

Arrayed assemblies were performed in an 96- or 384-position format by mating arrays of donor strains with an arrays of barcoded recipient strains, then pinning to selectable agar pads and counter-selectable agar pads in series using a Singer ROTOR HDA pinning robot. Recipient arrays were grown at 30°C overnight on LB+ 25 *μ*g/mL Gm+100 *μ*g/mL Carb, and the donor arrays were grown at 37°C overnight on LB+ 100 *μ*g/mL Hyg+ 2 *μ*g/mL Tc. The donor and recipient colonies were then pinned and mixed onto the same mating plate (LB+0.2% arabinose+0.2% rhamnose) and incubated at 30°C for 4-6 hours. To select for recombinant recipient cells, the mated colonies were pinned onto a selection plate LB+ 100 *μ*g/mL Hyg+ 25 *μ*g/mL Gm +100 *μ*g/mL Carb + 0.2% arabinose and incubated at 30°C overnight. The next day, cells on the selection plate were pinned to the counter-selection plate LB+ 100 *μ*g/mL Hyg+25 *μ*g/mL Gm +100 *μ*g/mL Carb + 200 *μ*g/mL 4CP and incubated at 30°C overnight, completing the first stitch. In parallel, donor cells for the second stitch were grown at 37°C overnight on either LB+60 *μ*g/mL Sp+ 2 *μ*g/mL Tc plate or LB+100 *μ*g/mL Apm+ 2 *μ*g/mL Tc plate.

To initiate the second stitch, recombinant recipients from the first stitch and donors for the second stitch were pinned and mixed onto the same mating plate (LB+0.2% arabinose+0.2% rhamnose) using a Singer ROTOR and incubated at 30°C for 4-6 hours. To select for only the recombinant recipient cells, the mixed cells were then pinned onto either an LB+ 60 *μ*g/mL Sp+ 25 *μ*g/mL Gm +100 *μ*g/mL Carb+0.2% arabinose plate or LB+100 *μ*g/mL Apm+ 25 *μ*g/mL Gm+100 *μ*g/mL Carb + 0.2% arabinose plate, depending on the the selection marker in the donor cells. After overnight growth at 30°C, cells on the selection plate were pinned to either an LB+60 *μ*g/mL Sp +25 *μ*g/mL Gm + 100 *μ*g/mL Carb + 6% sucrose plate or LB+ 100 *μ*g/mL Apm+ 25 *μ*g/mL Gm+100 *μ*g/mL Carb+6% sucrose plate and incubated at 30°C overnight, completing the second stitch. In parallel, donor cells for the third stitch were grown at 37°C overnight on LB+100 *μ*g/mL Hyg+2 *μ*g/mL Tc plate.

The assembly process and agar plates used for every odd stitch onwards (third, fifth, seventh, etc.) is the same as the first stitch, and the assembly process and agar plates used for every even stitch onwards (fourth, sixth, etc.) is the same as the second stitch. Following the last round of assembly, the helper plasmid with the temperature-sensitive origin was removed from the recipient cells by transferring the cells to an extra round of counter-selection plates without 100 *μ*g/mL Carb and growing them at 37°C overnight.

### Arrayed assembly in liquid

Arrayed assemblies were conducted in liquid in 96-well plates. The strains, media, and antibiotics used for each stitching step are the same as those in the agar assembly described above. To initiate the stitching process in a 96-well plate, the barcoded array of recipient strains and donor strain array were grown in 150 *μ*L of LB+25 *μ*g/mL Gm+100 *μ*g/mL Carb, and LB+100 *μ*g/mL Hyg+ 2 *μ*g/mL Tc, respectively, on an orbital plate shaker at 37°C overnight with 225 rpm shaking. Then, 120 *μ*L of the donor cells were mixed with 30 *μ*L of recipient cells and centrifuged. The supernatant was removed using a multi-channel pipette, and the cells were resuspended in 100 *μ*L of LB+0.2% arabinose+0.2% rhamnose. The mixed cells were mated for 4-6 hours at 30°C with 225 rpm shaking. To select for the recombinant recipient cells, 15 *μ*L of the mixed cells were transferred into 135 *μ*L of LB+100 *μ*g/mL Hyg+25 *μ*g/mL Gm+ 100 *μ*g/mL Carb selection media and grown overnight at 30°C with 225 rpm shaking. The next day, 5 *μ*L of the overnight culture was transferred into 145 *μ*L of LB+100 *μ*g/mL Hyg+ 25 *μ*g/mL Gm+100 *μ*g/mL Carb+200 *μ*g/mL 4CP counter-selection media and grown overnight at 30°C with 225 rpm shaking. In parallel, donor cells for the second stitch were grown at 37°C overnight with 225 rpm shaking in 150 *μ*L of either LB+60 *μ*g/mL Sp+ 2 *μ*g/mL Tc or LB+100 *μ*g/mL Apm+2 *μ*g/mL Tc.

To initiate the second stitch, 30 *μ*L of recombinant recipients from the first stitch and 120 *μ*L of donors for the second stitch were mixed and centrifuged. The supernatant was removed using a multi-channel pipette, and the cells were resuspended in 100 *μ*L of LB+0.2% arabinose+0.2% rhamnose. The mixed cells were mated for 4-6 hours at 30°C with 225 rpm shaking. To select for the recombinant recipient cells, 15 *μ*L of the mixed cells were transferred into 135 *μ*L of either LB+60 *μ*g/mL Sp+25 *μ*g/mL Gm+ 100 *μ*g/mL Carb or LB+100 *μ*g/mL Apm+ 25 *μ*g/mL Gm+100 *μ*g/mL Carb selection media and grown overnight at 30°C with 225 rpm shaking. The next day, 5 *μ*L of the overnight culture was transferred into 145 *μ*L of either LB+60 *μ*g/mL Sp+25 *μ*g/mL Gm+100 *μ*g/mL Carb+6% sucrose or LB+100 *μ*g/mL Apm+25 *μ*g/mL Gm+100 *μ*g/mL Carb+6%sucrose counter-selection media and grown overnight at 30°C with 225 rpm shaking. In parallel, donor cells for the third stitch were grown at 37°C overnight with 225 rpm shaking in 150 *μ*L of LB+100 *μ*g/mL Hyg+2 *μ*g/mL Tc.

The assembly process and media used for every odd stitch onwards (third, fifth, seventh, etc.) are the same as the first stitch, and the assembly process and media used for even stitch onwards (second, fourth, sixth, etc.) are the same as the second stitch. Following the last round of assembly, the helper plasmid with the temperature-sensitive origin was removed from the recipient cells by transferring 5 *μ*L of cells to an extra round of 145 *μ*L of counter-selection media without 100 *μ*g/mL Carb and growing them at 37°C overnight.

### Design of the GPCR variant library

We aimed to create a library of novel G protein-coupled receptor (GPCR) chimeras for high-throughput ligand screening in engineered yeast cells. Utilizing GPCRdb, we performed a multiple sequence alignment of 56 human GPCRs that are functional in yeast^20,26,27^. By applying a 15-amino acid (45-bp) percent identity sliding window average over the alignment and excluding positions with gaps, we identified three local maxima in transmembrane helices 2, 4, and 6. These local maxima served as fixed homology regions for our assemblies, resulting in the creation of four DNA blocks per gene.

To minimize mutations in the three 45-bp homology regions, gene selection was prioritized based on higher percent identity in these regions from the initial alignment. For future library use in yeast, we selected GPCRs with hydrophilic ligands, closely related family members, and those with available structural and pharmacological data. The chosen receptors included two adenosine receptors (ADORA2A, ADORA2B), three serotonin receptors (HTR1A, HTR1D, HTR4), and an epinephrine receptor (ADRB2) (**Supplementary Table 7**). The amino acid sequences were codon optimized for Saccharomyces cerevisiae using Integrated DNA Technology’s Codon Optimization Tool (https://www.idtdna.com/CodonOpt).

### Pooled assembly in liquid

We performed four independent pooled assemblies (two technical replicates of two biological replicates using different donor colones with the same DNA block sequences). Cell pools were grown in standard 50 mL conical tubes. For each stitch, 33 OD of donor cells (5.5 OD for each donor strain) and 8.25 OD of recipient cells from the previous stitch were mixed and spun down at 3,900 rpm for 10 minutes. The cell pellets were suspended in 10 mL LB+ 0.2% arabinose + 0.2% rhamnose medium and incubated at 30°C for 4-5 hours with 250 rpm shaking. Based on an estimated stitching efficiency of 10^−5^ calculated above, we expected ∼2.5*10^4^ stitching events in the pool (2.48*10^9^ cells/10^−5^) or ∼20X the number of designs. Cell cultures were spun down, resuspended in 30 mL of selection medium (first stitch and odd stitch selection: LB+200 *μ*g/mL Hyg + 25 *μ*g/mL Gm +100 *μ*g/mL Carb + 0.2% arabinose; even stitch selection: LB+ 60 *μ*g/mL Sp+ 25 *μ*g/mL Gm + 100 *μ*g/mL Carb + 0.2% arabinose), and incubated at 30°C overnight with 250 rpm shaking. Next, 15 mL of selected cells were spun down and resuspended with 45 mL of the same selection medium for 5 hours at 30°C with 250 rpm shaking. For counter-selection, 5 mL of of this culture was mixed with 45 mL counter selection medium (first stitch and odd stitch counter-selection: LB+200 *μ*g/mL Hyg+ 25 *μ*g/mL Gm +100 *μ*g/mL Carb + 200 *μ*g/mL 4CP; even stitch counter-selection: LB+ 60 *μ*g/mL Sp +25 *μ*g/mL Gm +100 *μ*g/mL Carb + 6% sucrose) at 30°C overnight with 250 rpm shaking. Donor cells for the following stitch were cultured on the same day at 37°C overnight with 250 rpm shaking. Following the last round of assembly, the helper plasmid with the temperature-sensitive origin was removed from the recipient cells by doing an extra round of counter-selection without 100 *μ*g/mL Carb and incubated at 37°C overnight.

### Data analysis for pooled GPCR assembly

We sequenced the pool of GPCR assembly products at every stitch by Oxford Nanopore Sequencing and mapped each read to the 1,296 GPCR designs using minimap2 (v2.28) using minimap2 -c options to generate paf file^28^. The paf files provided the mapped reference and the number of residue matches for each read. Reads that mapped to multiple GPCR variants were assigned to the assembly product with the highest number of residue matches. Reads that matched equally to two assembly products were not counted. **Supplementary Table 8** records the number of reads that mapped to each design before and after filtering. Some block combinations that were not detected by sequencing at early stages of the assembly were detected at a later stage of the assembly, indicating the sequencing data missed some designs present in the pool (**Supplementary Table 9**). The analysis results are summarized in **Supplementary Table 10**. Python package *muller* v0.6.0 is used for making Muller plots^29^.

### Sequence verification of single colonies from pooled GPCR assembly

Based on the frequency of stitching in liquid of 10^−5^, we expected that each assembly was the product of multiple assembly lineages, some of which may contain an error. Because of the high error rate of Oxford Nanopore sequencing (Q18-20), we could not determine if an error of given read was due to a sequencing error or a true error in a lineage. To more accurately determine the stitching fidelity of the pooled assembly, the four pooled GPCR assemblies were streaked onto LB+ 100 *μ*g/mL Sp+25 *μ*g/mL Gm agar plates and grown overnight at 37°C. From each replicate, 24 colonies were picked, and plasmids were extracted using the Qiagen MiniPrep kit. The plasmids were then individually sequenced (Plasmidsaurus/Primordium). Alignments (minimap2) between the consensus sequences generated by Plasmidsaurus/Primordium and the expected reference sequences were used to estimate the error rate (**Supplementary Table 11**).

### Examination of sequence features in constructs

For each construct assembled in this paper, several sequence features were examined: interspersed repeats, extreme GC content, and secondary structures. To search for interspersed repeats, BLAST (v2.16.0) “blastn” function was used to align the construct sequence to itself with the following parameters: -word_size 6, -reward 1, -penalty −1, -gapopen 3, -gapextend 2, -evalue 5, and -perc_identity 70. To identify regions with extremely high or low GC%, the GC% in a 51 bp rolling window was calculated. To find any secondary structures, Python package “Primer3” (primer3-py v2.0.3) was used with default parameters except for hairpin_window_size, which was changed to 41.

### Choosing regions of homology between DNA blocks

DNA blocks for each construct, along with their corresponding homology regions, were manually designed while taking into consideration specific sequence features that may affect homologous recombination efficiency and fidelity, as described above. For fluorophores, yeast chromosomes, and synthetic DNA, variable-sized DNA blocks were created to ensure that the homology regions had minimal repetitive sequences, homopolymers, and secondary structures, while maintaining a GC content close to 50%. For BGCs, all DNA blocks were ordered as 1,800 bp fragments. If homology regions contained repetitive sequences, homopolymers, secondary structures, or extreme GC content, the length of the homology region was extended up to 100 bp, depending on the extent of these sequence features.

### Oxford Nanopore Technologies (ONT) sequencing and data analysis

For high-throughput in-house sequencing, arrays of cells containing barcoded recipient plasmids (with barcodes marking position on a plate) were pooled by adding 10 mL of water to each agar plate and scraping the cells using a cell spreader. Since the plasmid size can vary significantly depending on the assembly (ranging from ∼6.5 kb to ∼27 kp) and size may impact the plasmid prep efficiency, we designed assembly plates such that all positions contained plasmids within a 1 kb range of each other. Any plates with overlapping barcodes or large differences in plasmid size were pooled separately. After pooling, the plasmids were extracted from the harvested cells using Qiagen Miniprep kits.

Sequencing libraries for each plasmid pool were prepared using the Rapid Barcoding Kit 96 V14 (SQK-RBK114.96) and run on Nanopore MinION Flow Cell (R10.4.1; FLO-MIN114). Each plasmid pool that was run on the same flow cell was uniquely barcoded with a different Rapid Barcode. The libraries were prepared using the standard protocol, with the following modifications: 1) 200 ng of input DNA was used instead of 50 ng, 2) 1.5 *μ*L of the Rapid Barcode reagent was used instead of 1 *μ*L, 3) DNA was eluted from the AMPure XP beads using 15 *μ*L of Elution Buffer EB per 24 Rapid Barcodes instead of a flat 15 *μ*L, and 4) up to 2 *μ*g of a DNA library was loaded onto the flow cell instead of the recommended maximum of 800 ng. For each sequencing run on a flow cell, we aimed for at least 100x coverage per plasmid. To minimize bias in sequencing due to plasmid size, only plasmid pools within a 5 kb range of each other were run together on the same flow cell. Data from the flow cell was basecalled using Dorado (https://github.com/nanoporetech/dorado) with Super-accuracy basecalling. The data were then demultiplexed by Rapid Barcodes using Dorado.

For runs with a small number of plasmids, the flow cells were stopped when the number of reads per rapid barcode reached ∼200 times the number of plasmids. The flow cells were then washed using the Flow Cell Wash Kit (EXP-WSH004) following the provided protocol before loading the next set of DNA libraries.

ONT reads were analyzed by custom developed software in Python and executed on a local 36-core cluster. The pipeline begins with a demultiplexing step where sequencing reads are separated based on custom barcodes in the recipient plasmid using a tailored demultiplexing program. Following demultiplexing, the reads are aligned to reference sequences with minimap2 (v2.28) using the “map-ont” option^28,30^. The alignment files are sorted and indexed using samtools (v1.20) using default parameters^31^. Variant calling is performed using bcftools^31^ (v1.20) with the *mpileup* (default ont-sup-1.20 parameters, except the coefficient for modeling homopolymer errors to 200, and max-depth to 10,000), and *call* (default parameters, except ploidy is set to 1) commands to identify small mutations and indels. Large structural variants (indels >10 bp) are identified with Sniffles2 (v2.2) using default parameters^32^.

To estimate the purity score, which represents the fraction of plasmid molecules with correctly assembled full-length products within an assembly replicate, read lengths were extracted from the demultiplexed FASTQ files. A histogram with a bin size of 50 bp was then created. “Peaks” or enrichment of certain read lengths, which presumably represents sequence data from fully intact plasmid DNA molecules that were not fragmented during plasmid extraction and library preparation, were identified using SciPy (v1.14.0) “find_peaks” function, with several changes from the default parameters. To account for the difference in read coverage, different values were used for the *height a*nd *prominence* parameters. The *height* parameter was set to 8, or to 5% of the number of reads in the largest bin (most number of reads), whichever is higher. The *prominence* parameter was set to 3.1, or two times the standard deviation in number of reads across all bins, whichever is higher. The *distance* parameter was set to two. After the peaks were detected, reads were assigned to the nearest peak to form clusters. Any reads farther than 150 bp away from a cluster peak (presumably caused by partial sequencing reads from fragmented DNA during library preparation) were not assigned to a cluster. The purity estimation of each sample was calculated by determining the proportion of reads clustered within 150 bp of the expected plasmid size over all reads assigned to any cluster. The reference mapping and variant calling analysis is reiterated on clustered reads to refine the variant calls and QC assessments. Only point mutations called with high confidence (combined quality score > 100) were output in the final QC report generated by the nanopore sequencing analysis pipeline. Any identified mutations and indels were categorized based on whether they were in the assembled product or the plasmid backbone.

We next adjusted the purity scores calculated above to only consider errors that impact the assembly fidelity. For the read cluster of the expected size, if any point mutations or indels were found in the assembly sequence, the purity score was assigned to zero, with the exception of point mutations that could be traced back to being present in the donor plasmids/cells. Mutations found in donor plasmids were ignored, since these mutations do not reflect assembly errors and are likely to stem from errors in synthesis. Mutations or deletions <100 bp that were detected in the plasmid backbone were also ignored. We rarely encountered instances where large structural rearrangements (indels >100 bp) in the plasmid backbone resulted in a lower purity score by forming a separate read length cluster, despite having a sequence-perfect assembly region. In these cases, we did not adjust the purity score, as the large indels could indicate plasmid instability and an undesirable plasmid product. Purity scores for all samples are found in **Supplementary Table 4**. All variants detected for all samples are in **Supplementary Table 5**.

The identities of insertions called by Sniffles2 (v2.2) (N=37) were mapped using minimap2 with default parameters against the sequences of the donor and recipient plasmids, and the sequences of the donor strain genome and F-plasmid.

For replicate assemblies of the three fluorophores (mPapaya, mPlum, sfGFP, 4 stitches each), we assumed a replicate contained no errors if >95% of molecules are sequence perfect. Errors in these assemblies were caused by a subpopulation of molecules with point mutations or undesired assembly products. For other assemblies performed here, we reported the purity score directly and discussed errors in detail in the Results section.

### GPCR protein structure

The predicted protein structure for a GPCR was generated in AlphaFold using the human GPCR variant *ADRB2*, which is one of the GPCRs in our test set (only it has been modified at the homology regions). The structure, colored by block boundaries, was generated using the Structure Viewer tool at: https://alphafold.ebi.ac.uk/entry/X5DQM5.

### Testing whether deletions occur more often in regions with long repeats

To assess whether deletions are more likely to occur in regions with repeated sequences, the 101 bp window surrounding the start and end positions of identified deletions was locally aligned to each other using BLAST (v2.16.0). If the same deletion was detected multiple times across different read length clusters within an assembly replicate, the deletion was only considered once. A total of 117 deletions total were examined. The BLAST “blastn” function was employed with the following parameters: -word_size 4 -reward 1 -penalty −1 -gapopen 5 -gapextend 2. For each deletion, 100 random pairs of 101 bp windows, separated by the same distance as the deletion, were randomly chosen from the construct where the deletion was detected. Each random pair of windows was also locally aligned to each other with blastn using the same parameters. A one-sided Wilcoxon rank-sum test was then used to compare the max BLAST scores with those of the randomly selected regions to test for significant differences.

### Construction of bridge donor plasmids

Bridge sequences that are 80-140 bp long were used to stitch together DNA blocks without homology. To scarlessly stitch full-length DNA blocks together, the first 40-70 bp of the bridge sequence was designed to match the last 40-70 bp of the preceding DNA block, and the second 40-70 bp was designed to match the first 40-70 bp of the next incoming DNA block. To stitch together parts of existing DNA blocks, the first 40-70 bp of the bridge sequence was designed to match the last 40-70 bp of the desired region in the preceding DNA block, and the second 40-70 bp was designed to match the first 40-70 bp of the desired region in the next incoming DNA block.

To construct the bridge donor plasmids, three different sets of bridge sequences, each flanked by AscI and NotI restriction sites, were designed on the same DNA fragment and ordered from TWIST. The TWIST DNA fragment was then digested using AscI and NotI restriction enzymes, and fragments of the expected size were gel extracted. The extracted gel fragment was ligated with AscI and NotI digested odd or even donor plasmids using T4 ligase and transformed into the dSL2 donor strain. Cells carrying the correct bridge donor plasmid were selected on LB+100 *μ*g/mL Hyg+2 *μ*g/mL Tc agar plates for odd donor plasmids, and with LB+60 *μ*g/mL Sp+2 *μ*g/mL Tc agar plates for even donor plasmids. The identity of the donor plasmids were then determined by Sanger sequence using primer oSL8 (**Supplementary Table 13**).

For stitching together two DNA blocks where one is in an odd donor plasmid or the other is in an even donor plasmid, only one stitch with a donor plasmid containing the bridge sequence is necessary (e.g., odd - even bridge - odd). However, for stitching together pairs of DNA blocks where both are in odd donor plasmids or even donor plasmids, two stitching steps are required to ensure that the selection and counter-selection markers are in the correct orientation (e.g., odd - even bridge 1 - odd bridge 2 - even). In these cases, the bridge sequence is the same for both odd and even bridge donor plasmids. Because the bridge sequence remains the same, the assembled product on the recipient remains the same; only the downstream selection and counter-selection cassettes are exchanged.

### Transfer of assembled products to donor plasmids

To transfer fully assembled products from recipient plasmids to donor plasmids, the recipient plasmids were digested with AscI and NotI restriction enzymes, and DNA fragments of the expected size were gel extracted. The gel extracted fragments were ligated with AscI and NotI digested donor plasmids using T4 ligase, and transformed into the dSL2 donor strain. Cells carrying the correct donor plasmids were selected on LB +100 *μ*g/mL Hyg+2 *μ*g/mL Tc agar plates.

## References

1. Gibson, D. G. et al. Enzymatic assembly of DNA molecules up to several hundred kilobases. Nat. Methods 6, 343–345 (2009).

2. Engler, C., Gruetzner, R., Kandzia, R. & Marillonnet, S. Golden Gate Shuffling: A One-Pot DNA Shuffling Method Based on Type IIs Restriction Enzymes. PLOS ONE 4, e5553 (2009).

3. Weber, E., Engler, C., Gruetzner, R., Werner, S. & Marillonnet, S. A Modular Cloning System for Standardized Assembly of Multigene Constructs. PLOS ONE 6, e16765 (2011).

4. Hillson, N. et al. Building a global alliance of biofoundries. Nat. Commun. 10, 2040 (2019).

5. Ma, Y., Zhang, Z., Jia, B. & Yuan, Y. Automated high-throughput DNA synthesis and assembly. Heliyon 10, e26967 (2024).

6. Li, M. Z. & Elledge, S. J. MAGIC, an *in vivo* genetic method for the rapid construction of recombinant DNA molecules. Nat. Genet. 37, 311–319 (2005).

7. Wang, K. et al. Defining synonymous codon compression schemes by genome recoding. Nature 539, 59–64 (2016).

8. Zürcher, J. F. et al. Continuous synthesis of E. coli genome sections and Mb-scale human DNA assembly. Nature 1–8 (2023) doi:10.1038/s41586-023-06268-1.

9. Jinek, M. et al. A programmable dual-RNA-guided DNA endonuclease in adaptive bacterial immunity. Science 337, 816–821 (2012).

10. Jiang, W., Bikard, D., Cox, D., Zhang, F. & Marraffini, L. A. RNA-guided editing of bacterial genomes using CRISPR-Cas systems. Nat. Biotechnol. 31, 233–239 (2013).

11. Thomason, L. C., Sawitzke, J. A., Li, X., Costantino, N. & Court, D. L. Recombineering: genetic engineering in bacteria using homologous recombination. Curr. Protoc. Mol. Biol. 106, 1.16.1–1.16.39 (2014).

12. Hoi, H. et al. An Engineered Monomeric Zoanthus sp. Yellow Fluorescent Protein. Chem. Biol. 20, 1296–1304 (2013).

13. Li, W. et al. Arrayed in vivo barcoding for multiplexed sequence verification of plasmid DNA and demultiplexing of pooled libraries. Nucleic Acids Res. 52, e47 (2024).

14. Oberortner, E., Cheng, J.-F., Hillson, N. J. & Deutsch, S. Streamlining the Design-to-Build Transition with Build-Optimization Software Tools. ACS Synth. Biol. 6, 485–496 (2017).

15. Chayot, R., Montagne, B., Mazel, D. & Ricchetti, M. An end-joining repair mechanism in Escherichia coli. Proc. Natl. Acad. Sci. 107, 2141–2146 (2010).

16. Levy, S. F. et al. Quantitative evolutionary dynamics using high-resolution lineage tracking. Nature 519, 181–186 (2015).

17. Bosch, J. From software product lines to software ecosystems. in Proceedings of the 13th International Software Product Line Conference 111–119 (Carnegie Mellon University, USA, 2009).

18. Oster, C. & Wade, J. Ecosystem requirements for composability and reuse: An investigation into ecosystem factors that support adoption of composable practices for engineering design. Syst. Eng. 16, 439–452 (2013).

19. Shetty, R. P., Endy, D. & Knight, T. F. Engineering BioBrick vectors from BioBrick parts. J. Biol. Eng. 2, 5 (2008).

20. Pándy-Szekeres, G. et al. GPCRdb in 2023: state-specific structure models using AlphaFold2 and new ligand resources. Nucleic Acids Res. 51, D395–D402 (2023).

21. Boyer, H. W. & Roulland-dussoix, D. A complementation analysis of the restriction and modification of DNA in *Escherichia coli*. J. Mol. Biol. 41, 459–472 (1969).

22. Jensen, S. I., Lennen, R. M., Herrgård, M. J. & Nielsen, A. T. Seven gene deletions in seven days: Fast generation of Escherichia coli strains tolerant to acetate and osmotic stress. Sci. Rep. 5, 17874 (2015).

23. Rakowski, S. A. & Filutowicz, M. Plasmid R6K Replication Control. Plasmid 69, 231–242 (2013).

24. Miyazaki, K. Molecular Engineering of a PheS Counterselection Marker for Improved Operating Efficiency in Escherichia Coli. BioTechniques 58, 86–88 (2015).

25. García-Nafría, J., Watson, J. F. & Greger, I. H. IVA cloning: A single-tube universal cloning system exploiting bacterial In Vivo Assembly. Sci. Rep. 6, 27459 (2016).

26. Lengger, B. & Jensen, M. K. Engineering G protein-coupled receptor signalling in yeast for biotechnological and medical purposes. FEMS Yeast Res. 20, foz087 (2020).

27. Kapolka, N. J. et al. DCyFIR: a high-throughput CRISPR platform for multiplexed G protein-coupled receptor profiling and ligand discovery. Proc. Natl. Acad. Sci. 117, 13117–13126 (2020).

28. Li, H. Minimap2: pairwise alignment for nucleotide sequences. Bioinformatics 34, 3094–3100 (2018).

29. Streck, A., Kaufmann, T. L. & Schwarz, R. F. SMITH: spatially constrained stochastic model for simulation of intra-tumour heterogeneity. Bioinformatics 39, btad102 (2023).

30. Li, H. New strategies to improve minimap2 alignment accuracy. Bioinformatics 37, 4572–4574 (2021).

31. Danecek, P. et al. Twelve years of SAMtools and BCFtools. GigaScience 10, giab008 (2021).

32. Smolka, M. et al. Detection of mosaic and population-level structural variants with Sniffles2. Nat. Biotechnol. 1–10 (2024) doi:10.1038/s41587-023-02024-y.

